# MDFIC2 is a sensory neuron-specific PIEZO channel auxiliary subunit

**DOI:** 10.1101/2025.10.26.684595

**Authors:** Zijing Zhou, Fei Dai, Delfine Cheng, Xiaonuo Ma, Seyedeh Farzaneh Omidkhoda, Jack Clarke, Huijing Zhang, Michael Laden, Yang Guo, Jinyuan Vero Li, Renjing Liu, Emily S Wong, Yixiao Zhang, Charles D Cox

**Author notes:** These authors contributed equally (Zijing Zhou, Fei Dai).

## Abstract

PIEZO channels are critical for sensory mechanotransduction. While MyoD-family inhibitor proteins were identified as PIEZO1 auxiliary subunits, their broader regulatory roles, particularly in sensory cells, remained unclear. Here we demonstrate native MDFIC and MDFI regulate endogenous PIEZO channel currents in various non-sensory cell types. However, neither MDFIC nor MDFI are expressed in primary sensory neurons. In these cell types we identified an uncharacterised member of this family, *Mdfic2*/*Gm765*, that shares the ability to physically bind to PIEZO1 and PIEZO2. MDFIC2 is selectively expressed in subsets of mechanosensitive neurons, including dorsal root ganglia, trigeminal ganglia, and vagal sensory neurons. Like its paralogues, MDFIC2 alters PIEZO1/2 mechanosensitivity and inactivation kinetics, converting them into high-threshold slowly inactivating mechanoreceptors. Extensive cryo-EM reveals a conserved binding pocket for these auxiliary subunits in the pore modules of both PIEZO1 and PIEZO2 mediated by the post-translationally modified distal C-termini of MyoD-family inhibitor proteins. This provides a comprehensive structural and functional characterisation of MyoD-family inhibitor proteins as PIEZO1/2 channel regulators and offers new insights into sensory physiology and mechanical pain mechanisms.

## Introduction

Since their discovery, mechanosensitive PIEZO ion channels have emerged as essential molecular sensors that translate mechanical stimuli into cellular responses across both sensory and non-sensory systems (1-4). Their role includes the sensing of light touch. (3), proprioception (5) and mechanical pain related signals (4, 6-9). PIEZO1 is broadly expressed in non-neuronal cell types including endothelial, epithelial, mesenchymal, and muscle cells, where it contributes to diverse physiological processes (10). In contrast, PIEZO2 expression is largely confined to sensory tissues, with particularly high levels in dorsal root ganglion (DRG) neurons, vagal sensory neurons of the nodose and petrosal ganglia, trigeminal ganglia, and Merkel cells (11).

In primary non-sensory cells across nearly all lineages, PIEZO channel currents consistently exhibit slower inactivation kinetics compared to those recorded in heterologous expression systems (12-20). This phenomenon is also observed in sensory cells; for example, dorsal root ganglion (DRG) neurons display distinct mechanosensitive currents characterized by slower inactivation. Mechanosensitive DRG populations are in fact classified by their inactivation profiles, defined as either rapidly adapting (RA), intermediately adapting (IA) or slowly adapting (SA). These currents could reflect both the presence of novel ion channels with different kinetics to PIEZO channels and the influence of yet unidentified regulatory factors that modulate PIEZO1/2 function. Understanding these mechanisms is critical for decoding the molecular basis of mechanosensation in sensory physiology.

The prevailing view of PIEZO2 activity in DRGs is that PIEZO2 is solely responsible for rapidly adapting currents, with its loss in mice leaving other current types such as IA and SA currents largely intact (3, 18, 21). In some subsets of DRGs, PIEZO1 is also expressed and the combined ablation of PIEZO1 and PIEZO2 reduces all current types in these cells (22), whereas over expression of PIEZO1 boosts IA/SA currents (21). In duck trigeminal neurons PIEZO2 siRNA reduced the peak currents of all kinetic types, suggesting that in some sensory cells PIEZO2 can contribute to slower adapting currents (23). This is mirrored in certain subsets of mouse DRGs (6) and in touch sensitive Merkel cells where slowly inactivating currents (∼10% of peak currents) are also reduced after *Piezo2* knockout (24). However, the mechanism(s) by which PIEZO1 or PIEZO2 could contribute to these slower kinetic components in sensory cells is unknown.

We recently identified a family of PIEZO channel auxiliary subunits through affinity capture mass spectrometry (25). We found that the MyoD-family inhibitor proteins MDFIC and MDFI significantly reduced PIEZO1/2 channel inactivation and increased activation threshold (25). Here, we identify and characterize a novel MyoD-family inhibitor gene, *Mdfic2*. This gene is abundantly expressed in specific subsets of dorsal root ganglion (DRG) neurons, trigeminal ganglion neurons and vagal sensory neurons. Like MDFIC and MDFI, MDFIC2 modulates the mechanosensitivity and slows the inactivation of both PIEZO1 and PIEZO2. Furthermore, the electrophysiological properties of the PIEZO2:MDFIC2 complex is tuneable by the PIEZO sensitizing molecule, STOML3. Using cryo-electron microscopy, we revealed a previously uncharacterized but conserved binding site within the pore domain of human PIEZO2, shared across this family of auxiliary subunits. This provides a complete structural and functional characterization of this family of auxiliary subunits where their tissue and cell-specific expression likely contributes to the important physiological functions of PIEZO channels.

## Results

### MDFIC/MDFI regulate endogenous PIEZO currents

To probe the ubiquity of auxiliary subunit regulation of PIEZO channels we surveyed native mechanically evoked currents in six human and murine cell lines that span a variety of developmental origins. This includes cells of mesenchymal origin (pre-adipocytes; 3T3-L1), atrial cardiomyocyte-like cells (HL-1), a sensory Merkel cell line (MCC13), epithelial cells (HeLa), oligodendrocyte-like cells (HOG) and primary isolated mouse lung endothelial cells. We treated these cells with either control non-targeting siRNA or a composite of *Mdfic* and *Mdfi* siRNA and then indented the cells using a polished glass rod while recording currents in whole-cell mode. In all control siRNA treated cells we identified two components: a rapidly inactivating component and a slowly/non-inactivating component consistent with PIEZO channels bound to MDFIC or MDFI (25-27)(**Fig. 1A-O**). In all cell lines, knocking down MDFIC and MDFI resulted in a reduced current at the end of the mechanical stimulus (I_remaining_). The same outcome was observed in native mouse lung endothelial cells (**Fig. 1P-R**). Thus, in all cells tested, MDFIC and/or MDFI regulated native mechanosensitive channel inactivation (**Fig. 1**). As expected, the combined knockdown of PIEZO1 and PIEZO2 eliminated almost all native poking-evoked currents in all cell lines. These data combined indicate widespread use of MDFIC/MDFI as regulators of PIEZO channel function across diverse cell types.

**Figure 1.**
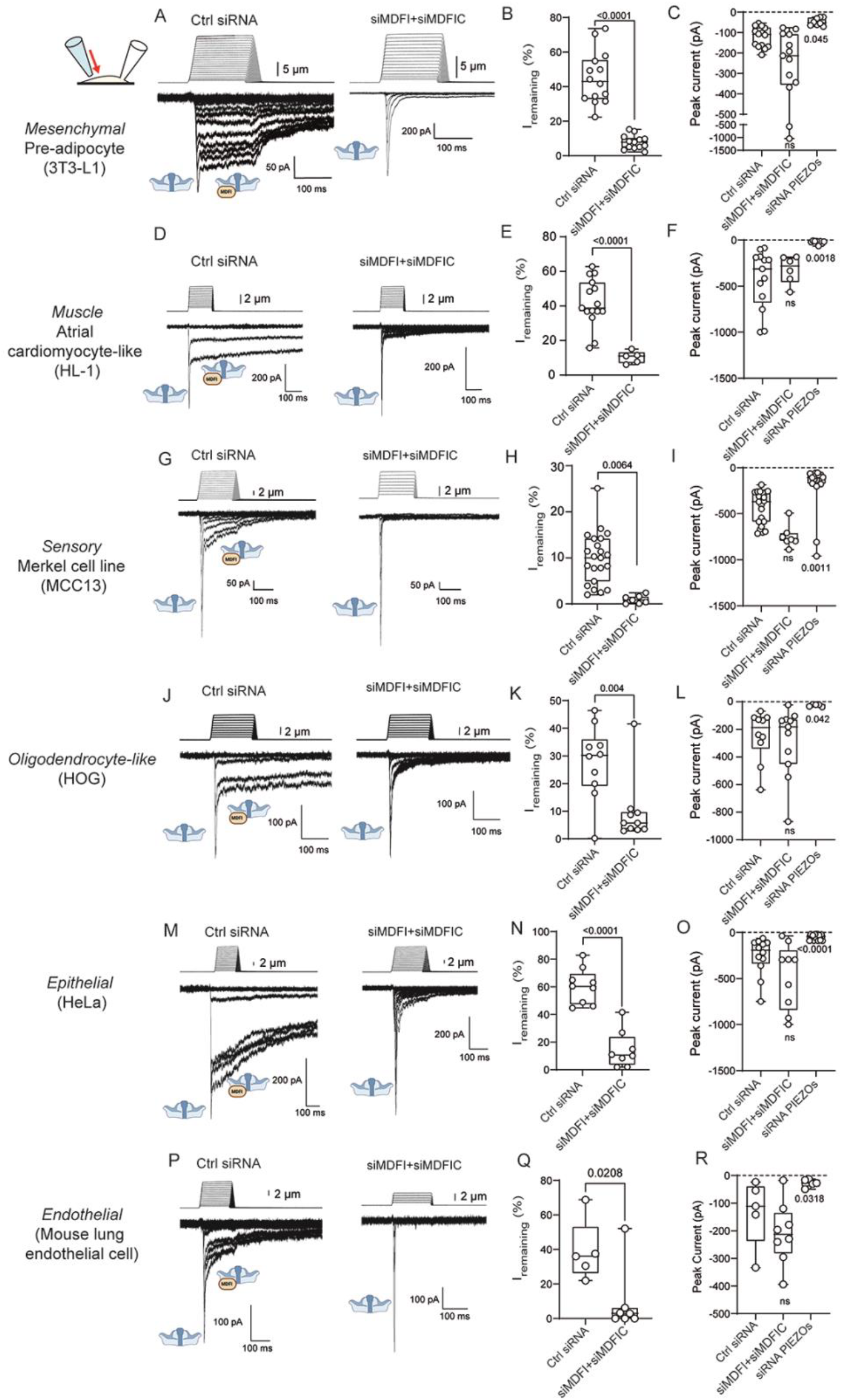
MDFIC and MDFI regulate native PIEZO currents in cell types from multiple lineages. Representative whole-cell traces of poking-evoked currents (two left panels), current remaining at the end of the poking stimulus (third panel) and peak currents elicited prior to cell rupture (right panel) in cells treated with control siRNA, siRNA for MDFIC and MDFI or siRNA for Piezo1 and Piezo2 in (A-C) 3T3-L1 cells, (D-F) HL-1 cells, (G-I) Merkel MCC13 cells, (J-L) human oligodendroma (HOG) cells, (M-O) HeLa cells, and (P-R) mouse lung endothelial cells. (Data is plotted as min to max box and whiskers plots; p-values were determined either using Student’s t-test for I_remaining_ plots or one-way ANOVA with Tukey’s multiple comparison for peak current plots).

### A new uncharacterized auxiliary subunit for sensory PIEZOs

We next asked if we could identify a primary sensory cell type that expressed MyoD-family inhibitor proteins. We focussed first on DRG neurons, given the multitude of mechanosensitive channel kinetic types present in these cells (28). However, analysis of published single cell transcriptomes of mouse DRGs (29, 30) revealed that there was little to no expression of *Mdfic* or *Mdfi* in any DRG populations (**Supplementary Fig. 1**).

We searched the human proteome for proteins that harbour a sequence related to the conserved C-terminal PIEZO1-binding domain of MDFIC (25) that could be expressed in primary sensory cells. This search yielded the gene *Gm765* (labelled “DRG protein short isoform”), which encodes an uncharacterized 189-residue protein that shows significant homology to MDFIC and MDFI (**Supplementary Fig. 2**). Strikingly, interrogation of the single-cell RNAseq datasets described above revealed that *Gm765/Mdfic2* is abundantly and selectively expressed in subsets of non-peptidergic (NP) neurons (**Fig. 2A**). These are small diameter DRGs expressing *Mrgprd* (**Fig. 2B**) which likely act as nociceptors. In this setting *Mrgprd*+ DRGs are known to display intermediately and slowly adapting mechanically-activated currents that are reduced on the conditional knockout of PIEZO2 (6) and require larger forces for mechano-activation (31). Previous work has also identified *Gm765*/*Mdfic2* as a marker gene of these NP subpopulations (32).

**Figure 2.**
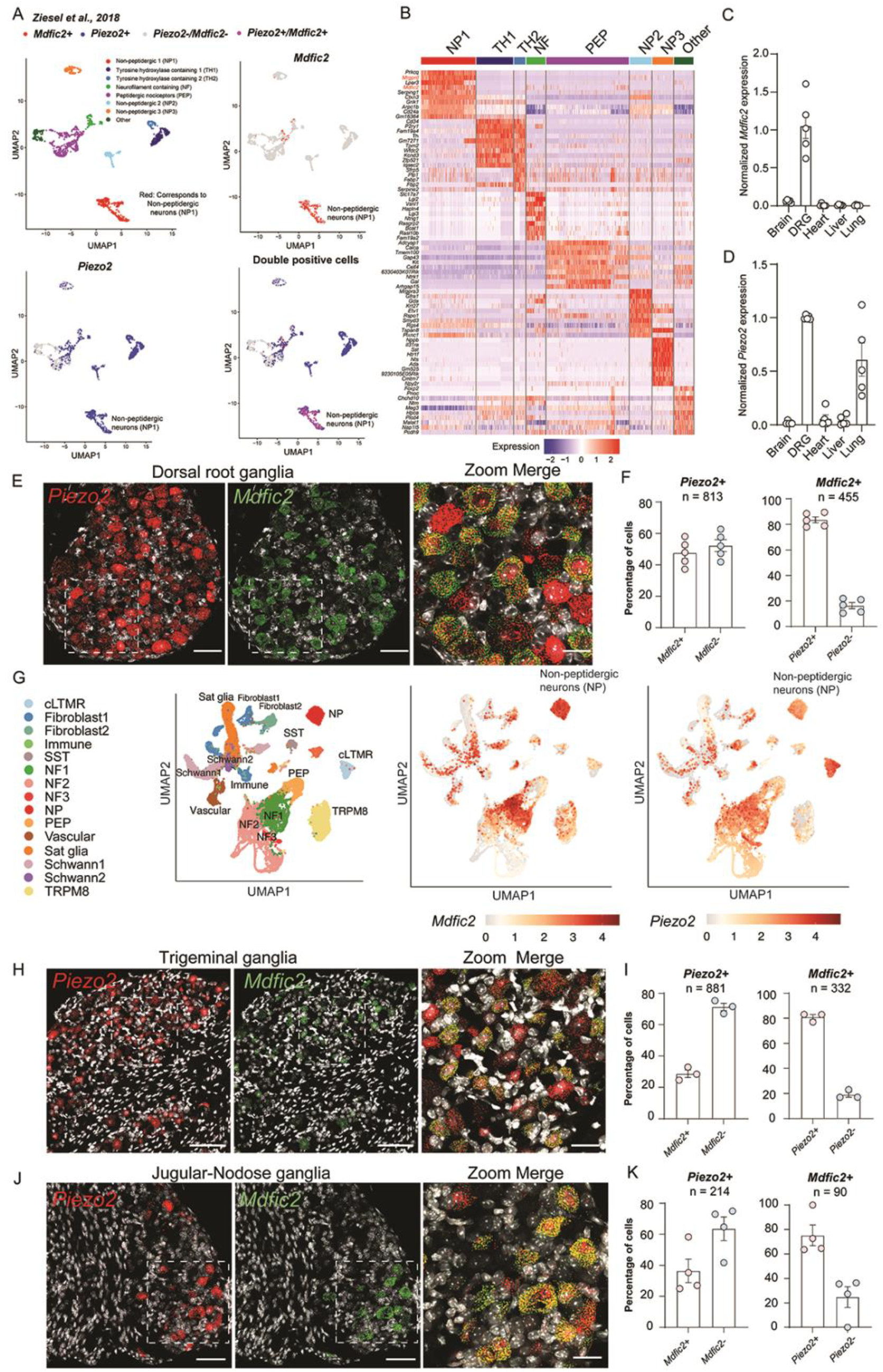
Mdfic2 is selectively expressed in subpopulations of dorsal root ganglion neurons, trigeminal ganglion neurons and vagal sensory neurons. (A) Single cell RNAseq from Ziesel et al., 2018 showing Mdfic2 and Piezo2 positive cells co-expressed in non-peptidergic neurons. (B) Differentially expressed genes in cell types from Ziesel et al., 2018 showing the non-peptidergic neurons expressing Mdfic2 are Mrgrpd+. Values reflect Z-score transformed log-normalized counts. (C-D) qPCR data from different tissues showing the high expression of MDFIC2 and PIEZO2 in DRGs. (E) RNAScope data showing Mdfic2 and Piezo2 RNA in a subset of DRG neurons. Nuclei are coloured white. Scale bars = 50 µm and 20 µm (zoomed image). (F) Statistics showing percentage of cells expressing both Mdfic2 and Piezo2, from 5 individual DRGs from 3 different animals. (G) ScRNAseq showing the expression of MDFIC2 in trigeminal ganglia in cell types transcriptionally similar to non-peptidergic neurons. (H) RNAScope data showing Mdfic2 and Piezo2 RNA present in a subset of TG neurons. Nuclei are coloured white. Scale bars = 50 µm and 20 µm (zoomed image). (I) Statistics showing percentage of cells expressing both Mdfic2 and Piezo2, from 3 individual TGs from 3 different animals. (J) RNAScope data showing Mdfic2 and Piezo2 RNA present in a subset of JNG neurons. Nuclei are coloured white. Scale bars = 50 µm and 20 µm (zoomed image). (K) Statistics showing percentage of cells expressing both Mdfic2 and Piezo2, from 4 individual JNGs from 3 different animals.

We isolated mouse DRGs and assessed *Mdfic2* expression by qRT-PCR and in four other organ tissues. *Mdfic2* was highly expressed in DRGs compared to other tissues, like the expression of *Piezo2*, with the highest detectable levels outside the DRGs being found in the brain (though expression in the brain was only ∼10% of that observed in DRGs; **Fig. 2C-D**). To visualize the expression pattern of *Piezo2* and *Mdfic2* we turned to RNAScope as we lacked reliable antibodies for both targets. Our data showed that 48% of *Piezo2* positive cells bear *Mdfic2* expression, while 84% of *Mdfic2* positive cells were *Piezo2* positive (**Fig. 2E-F**). We further found *Mdfic2* was expressed in a subset of DRGs which labelled for isolectin B4 (IB4), a marker of non-peptidergic DRG neurons (**Supplementary Fig. 3**). Consistent with the RNAseq we did not see any RNA signal in supporting cells that surround the DRG neurons (**Supplementary Fig. 3**). We also surveyed data from a single cell patchseq study of DRG neurons which showed that 6/53 cells expressed high levels of *Mdfic2* and that all six of these cells displayed slowly adapting mechanically-activated currents (28). We also noted that *Mdfic2* is downregulated in DRGs in models of neuropathic pain including the spared nerve injury (SNI) model (33) and chemotherapy-induced neuropathy (34).

In addition to DRGs *Mdfic2* is also expressed widely in human and mouse trigeminal ganglia (TG) including in cells transcriptionally similar to non-peptidergic DRG neurons that express *Mrgprd* (**Fig. 2G**) (35, 36). *Mdfic2* is also a marker gene of a subset of vagal sensory neurons in the jugular-nodose ganglia (JNG) (37, 38) that co-express PIEZO2 and innervate the lung and oesophagus and represent baroreceptors (a sensory cell in which both PIEZO1 and PIEZO2 are important for blood pressure sensing (39)). Our RNAScope data from *Mdfic2* and *Piezo2* on isolated mouse TGs and JNGs showed that *Mdfic2* was present in 27% and 36% of *Piezo2* positive cells, respectively (**Fig 2. H-K**).

Given the interesting and selective expression pattern of *Mdfic2* across sensory cell types and the significant homology to MDFIC and MDFI we cloned human *Mdfic2* with an N-terminal haemagluttinin (HA) tag and expressed it in *Piezo1*^*-/-*^ HEK293T cells alongside human PIEZO1 and PIEZO2. Using co-immunoprecipitation we showed that hMDFIC2 interacts with both PIEZO1 and PIEZO2 (**Supplementary Fig. 4A-B**) and using native gels we showed that MDFIC2 can be identified in a native complex with PIEZO1 (**Supplementary Fig. 4C**).

We next sought to assess whether MDFIC2 could regulate PIEZO currents in *Piezo1*^*-/-*^ HEK293T cells. After ∼72 hours of expression, we first surveyed the influence of MDFIC2 on stretch-evoked PIEZO1 currents and compared it to MDFIC (**Supplementary Fig. 5**). MDFIC2 slowed the activation of PIEZO1 in response to membrane stretch and completely ablated inactivation as represented by an increase in I_remaining_ from 13 ± 2.3 % in PIEZO1 only patches to 76.7 ± 4.8 % and 85.6 ± 1.8 % in the presence of MDFIC2 and MDFIC, respectively. The effect on currents after the pressure was released (I_post_) was less significant for MDFIC2 than for MDFIC (**Supplementary Fig. 5**).

To compare the effects of the three auxiliary subunits we expressed all three separately in wild-type (WT) Neuro2A (N2A) cells and characterized the response of the native PIEZO1 to indentation in whole-cell mode. In WT N2A cells, we observed that both MDFIC and MDFIC2 were recruited to the membrane (by colocalisation with the membrane marker LCK10-GFP) but displayed a distinct intracellular localization in *Piezo1*^*-/-*^ N2A cells (**Fig. 3A-B**). On expression of each of the MyoD-family inhibitor proteins we observed a marked slowing of channel inactivation compared to vector only transfected cells (**Fig. 3C**). It was also noticeable that while indenting N2A cells co-expressing MyoD-family inhibitor proteins that fewer cells responded with mechanically evoked PIEZO1 currents before patch rupture compared to vector only transfected cells. That is, 32/32 WT N2A cells responded, whereas only 16/22, 12/22, or 17/32 responded in the presence of MDFI, MDFIC or MDFIC2, respectively (**Fig. 3D**). Overall, the data showed a dramatic reduction in peak currents from all N2A cells expressing MyoD-family inhibitor proteins (**Fig. 3E**). The apparent threshold for activation also increased in the presence of MyoD-family inhibitor proteins as measured by the indentation depth at which currents exceeded 40 pA (**Fig. 3F**). This is consistent with previous work showing MDFIC increases the threshold for PIEZO1 activation (25). In addition to this change in mechanosensitivity there was a profound slowing of inactivation reminiscent of slowly adapting currents in DRG neurons (**Fig. 3G**).

**Figure 3.**
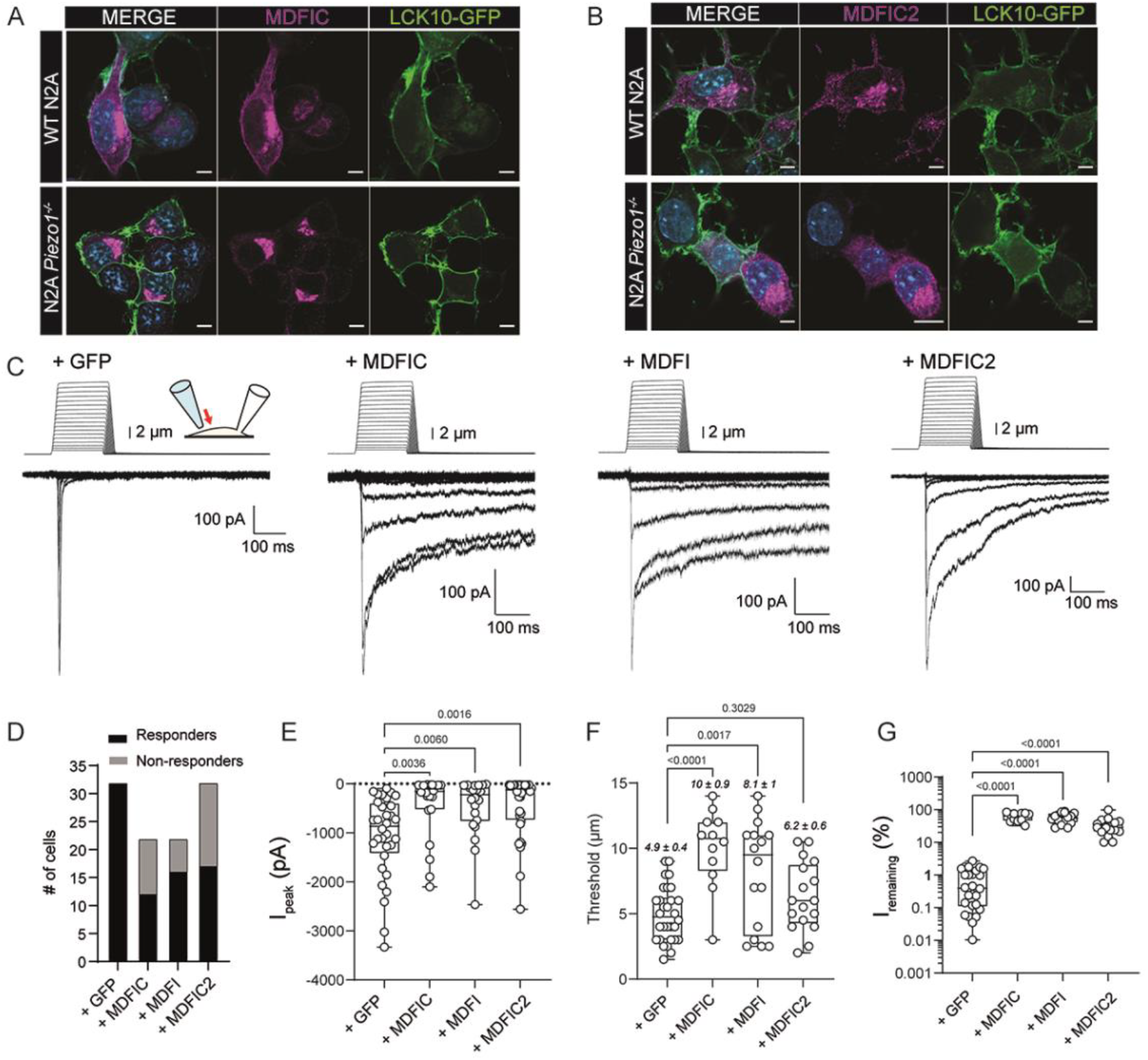
MDFIC2 and all MyoD-family inhibitor proteins regulate the inactivation kinetics and mechanosensitivity of native PIEZO1 currents in N2A cells. (A-B) Confocal images of N2A and Piezo1^-/-^ N2A expressing (A) hMDFIC and (B) hMDFIC2 alongside the membrane marker LCK10-GFP (scale bar = 10 µm). (C) Representative whole-cell traces of native N2A PIEZO1 currents at −50 mV in the presence of a vector only control and MDFIC, MDFI and MDFIC2. (D) Quantification of the number of cells responding to indentation from replicate recording of those shown in C. (E-G) Quantification of the peak current (E), threshold for activation of currents by indentation (F) and I_remaining_ at the end of the poke stimulus (G) for replicates of whole-cell recordings shown in C in the presence and absence of MyoD-family inhibitor proteins. (Data is plotted as min to max box and whiskers plots; p-values shown determined using one way ANOVA with Dunnett’s multiple comparison).

Using RNAScope we noticed a small population of cells (∼16% in DRG; ∼19% in ∼TG; ∼25% in JNG) that only expressed *Mdfic2* but not *Piezo2* perhaps indicative of an alternative PIEZO-independent role of MDFIC2 in sensory neurons. Given MDFIC binds to transcription factors such as GATA2 (40, 41) we tested binding of MDFI and MDFIC2 to this transcription factor using co-immunoprecipitation and found that all MyoD-family inhibitor proteins physically interact with GATA2 (**Supplementary Fig. 6)** suggesting a possible transcriptional regulatory role of MDFIC2 in sensory neurons.

### MDFIC2 converts PIEZO2 into a high threshold mechanoreceptor

Given PIEZO2 is more widely expressed in sensory neurons such as DRGs, we next addressed whether MDFIC2 could regulate PIEZO2 channel properties. We first investigated whether MDFIC2 could regulate human PIEZO2 in response to membrane stretch (**Fig. 4A-D**). While many reports suggested PIEZO2 was not as robustly activated by stretch as PIEZO1 we and others have recorded large and reproducible stretch activated currents (25, 42) using a plasmid encoding human PIEZO2 (43). After ∼72 hours expression in *Piezo1*^*-/-*^ HEK293T cells we saw a slowing of activation of hPIEZO2 in the presence of hMDFIC2 (**Fig. 4A-B**). We also observed an increase in the peak currents elicited per patch (**Fig. 4C**) and a slowing of inactivation indicated by an increase in I_remaining_ at the end of the pressure pulse (**Fig. 4D**). We also measured a right shift in the pressure response curve of human PIEZO2 in the presence of hMDFIC2 indicative of a reduced sensitivity of PIEZO2 to mechanical force (**Fig. 4E**). This is consistent with the effects in the cell-attached (25) and whole-cell configuration (**Fig. 3D,F**) observed for PIEZO1. We then surveyed the influence of MDFIC2 on PIEZO2 in whole-cell mode. We saw that again MDFIC2 slowed inactivation albeit not as significantly as seen for PIEZO1 (**Fig. 4F**). Here if we were to classify currents, the currents transitioned from rapidly adapting to intermediately adapting. However, all currents were best fit with a bi-exponential function where the expression of MDFIC2 increased the magnitude and contribution of the slower component (**Fig. 4G-H**) and the I_remaining_ (**Fig. 4I**). This was all consistent with MDFIC2 slowing PIEZO2 inactivation and the amount of MDFIC2 expression determining the degree of slowing. Corroborating the cell-attached data (**Fig. 4E**) we also observed clear and significant increase in the threshold for activation of PIEZO2 by indentation from 2.5 ± 0.3 µm to 5.3 ± 0.6 µm in the presence of MDFIC2 (**Fig. 4J**). These data indicate an MDFIC2-dependent reduction in the sensitivity of PIEZO2 to mechanical force.

**Figure 4.**
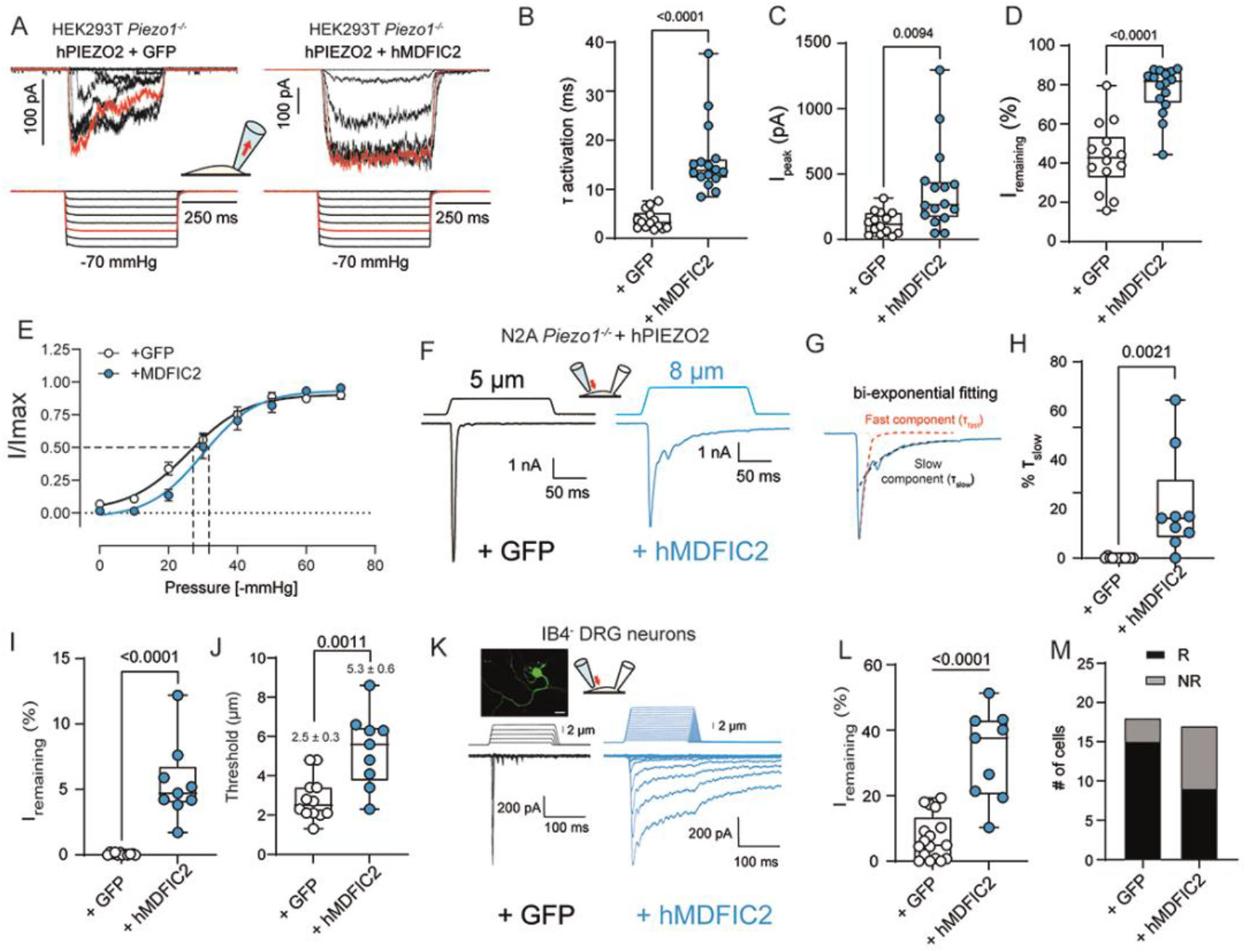
MDFIC2 regulates the mechanosensitivity and inactivation of human and mouse PIEZO2 currents. (A) Representative cell-attached traces of human PIEZO2 expressed in Piezo1^-/-^ HEK293T for ∼72 hours in the absence (left) and the presence (right) of hMDFIC2. Currents were elicited in response to negative pressure applied via a high-speed pressure clamp at a holding potential of −50 mV. (B-D) Quantification of time constant for activation, peak currents per patch and I_remaining_ for replicates of cell-attached recordings shown in A in the presence and absence of hMDFIC2. (E) Pressure response curves from data displayed in panel A showing a right shift of hPIEZO2:hMDFIC2 in the cell-attached configuration compared to hPIEZO2 alone. (F) Representative whole-cell traces of human PIEZO2 expressed in Piezo1^-/-^ N2A cells for ∼72 hours in the absence (left) and the presence (right) of hMDFIC2. (G-H) Whole-cell currents were analysed using biexponential fitting (G) and the contribution of the fast and slow components was quantified (H). (I-J) Quantification of I_remaining_ at the end of the poke stimulus and threshold for activation of currents by indentation for replicates of whole-cell recordings shown in F in the presence and absence of hMDFIC2. (K) Representative whole-cell traces from IB4 negative (IB4^-^) DRG neurons in the absence (left) and the presence of hMDFIC2 (right) at a holding potential of −50 mV. (L-M) Quantification of replicate recording of those shown in panel K measuring the current remaining at the end of the indentation stimulus (L) and the number of cells responding to a mechanical stimulus (M) (R – responders; NR – non-responders). (Data is plotted as min to max box and whiskers plots; p-values shown determined using t-test).

To test whether the degree of inactivation slowing was related to the relative levels of PIEZO2 compared to MDFIC2 we turned to primary isolated DRG neurons (**Fig. 4K-M**). In an attempt to maximize the saturation of PIEZO2 with MDFIC2, we electroporated DRGs with a plasmid encoding either GFP alone or MDFIC2 IRES GFP and compared the current kinetics in IB4 negative large diameter DRG neurons, which typically give rapidly adapting currents. In this scenario we saw a profound change in inactivation of mechanically evoked currents towards a slowly adapting phenotype. This is consistent with the idea that the ratio of PIEZO2 to MDFIC2 determines the absolute influence of the latter on inactivation kinetics. In addition, we observed a significant increase in the proportion of cells not responding to mechanical force (as we observed above for native PIEZO1 in N2A cells), reinforcing our conclusion that MDFIC2 reduces the sensitivity of PIEZO channels to a mechanical stimulus. These biophysical properties of the PIEZO2:MDFIC2 complex are more consistent with a high threshold mechanoreceptor.

### STOML3 sensitizes PIEZO2:MDFIC2 complexes

Conversely to the DRG setting, when co-expressed *in vitro*, MDFIC2 induced only a moderate alteration in the inactivation kinetics of PIEZO2. We reasoned that this may be due to the high threshold of MDFIC2 bound PIEZO2 channels, resulting in a large proportion of currents originating from apo-PIEZO2. If so, molecules known to sensitize PIEZO2 might amplify the observed effects on kinetics. To test this hypothesis, we co-expressed STOML3, a well-established sensitizer of PIEZO channels(44), together with PIEZO2 and MDFIC2 in *Piezo1*^*−/−*^ N2A cells and performed whole-cell electrophysiological recordings (**Fig. 5A**). Cells co-expressing MDFIC2 and STOML3 exhibited a marked left-shift in the mechanically evoked PIEZO2 currents compared to those expressing MDFIC2 alone, suggesting that STOML3 enhances the sensitivity of the PIEZO2:MDFIC2 complex (**Fig. 5B**). This was further supported by a significant reduction in the mechanical gating threshold upon STOML3 co-expression (**Fig. 5C**). While STOML3 alone did not alter the inactivation kinetics of PIEZO2, its sensitization effect facilitated the appearance of slowly inactivating currents in PIEZO2:MDFIC2 expressing cells, resulting in a significantly increased current remaining (**Fig. 5D**) akin to what we observed in the MDFIC2 overexpressed DRGs (**Fig. 4K**).

**Figure 5.**
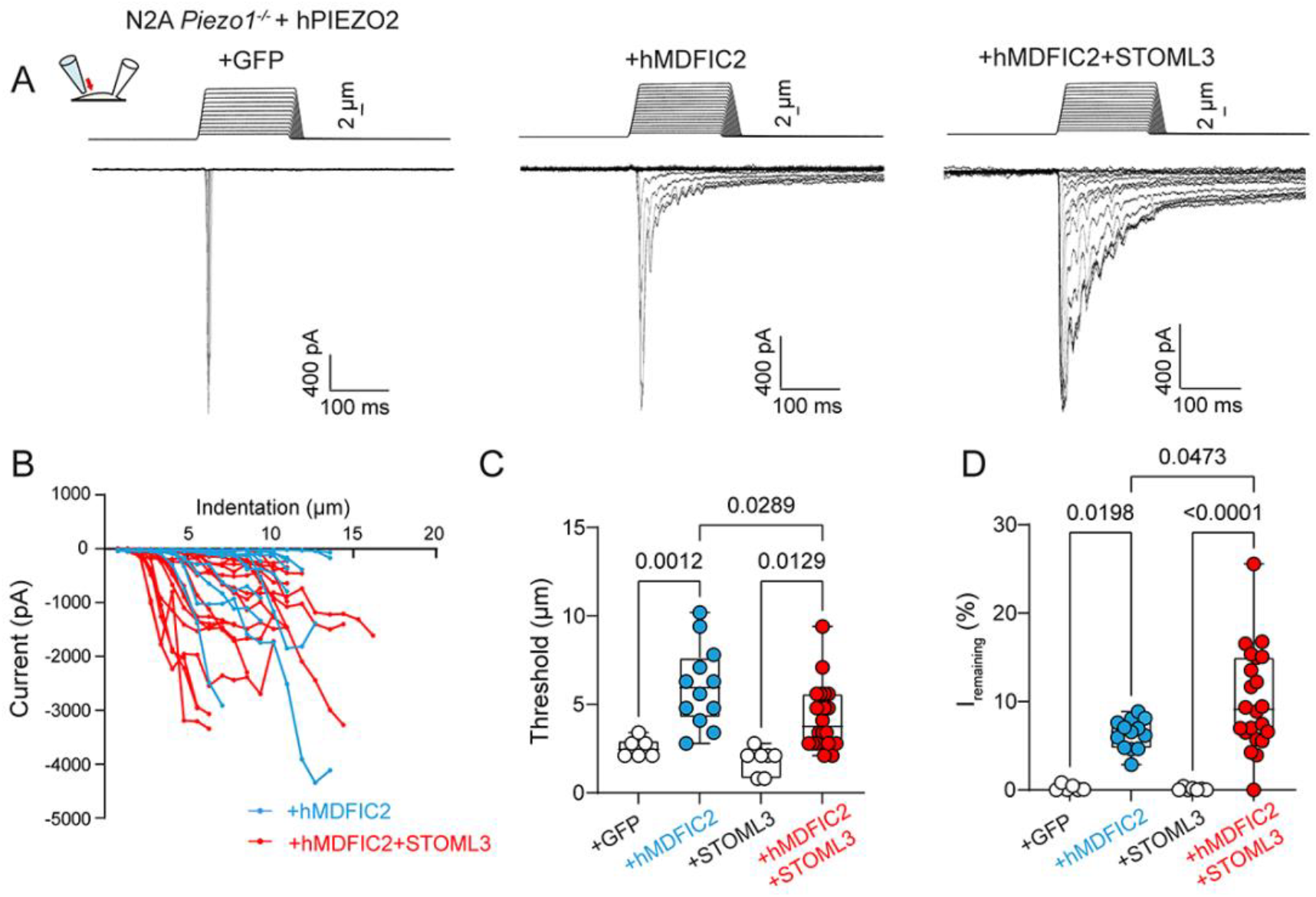
STOML3 can sensitize the PIEZO2:MDFIC2 complex. (A) Representative whole-cell traces of human PIEZO2 expressed in Piezo1^-/-^ N2A cells with GFP (left), hMIDFC2 (middle) or hMDFIC2/STOML3 co-expressed (right). (B) Whole-cell currents from individual cells were plotted against indentation depth. (C-D) Quantification of threshold of activation of currents by indentation (C) and I_remaining_ (D) at the end of the mechanical stimulus for replicates of whole-cell recordings shown in panel A. (Data is plotted as min to max box and whiskers plots; p-values shown determined using one-way ANOVA and Tukey’s multiple comparison test).

### MDFIC2 intercalates into the pore module of PIEZO2

To reveal the molecular details of the interaction of MDFIC2 with human PIEZO2 we turned to cryo-electron microscopy. Here we solved a suite of structures with examples of all MyoD-family inhibitor proteins bound to PIEZO channels. In all four of these structures, the PIEZO proteins had folds that closely resembled the previous structures of PIEZO1/2 (**Fig. 6; Supplementary Fig. 6-7; Supplementary Table 1**). In all cases, we observed three copies of the C-terminal helix of the MyoD-family inhibitor proteins bound to the threefold symmetry-related PIEZO-PIEZO interfaces. Specifically, this included mouse MDFI bound to mouse PIEZO1 (3.6 Å) and most importantly human MDFIC and human MDFIC2 bound to PIEZO2 at overall resolutions of 3.7 and 3.3 Å, respectively.

**Figure 6.**
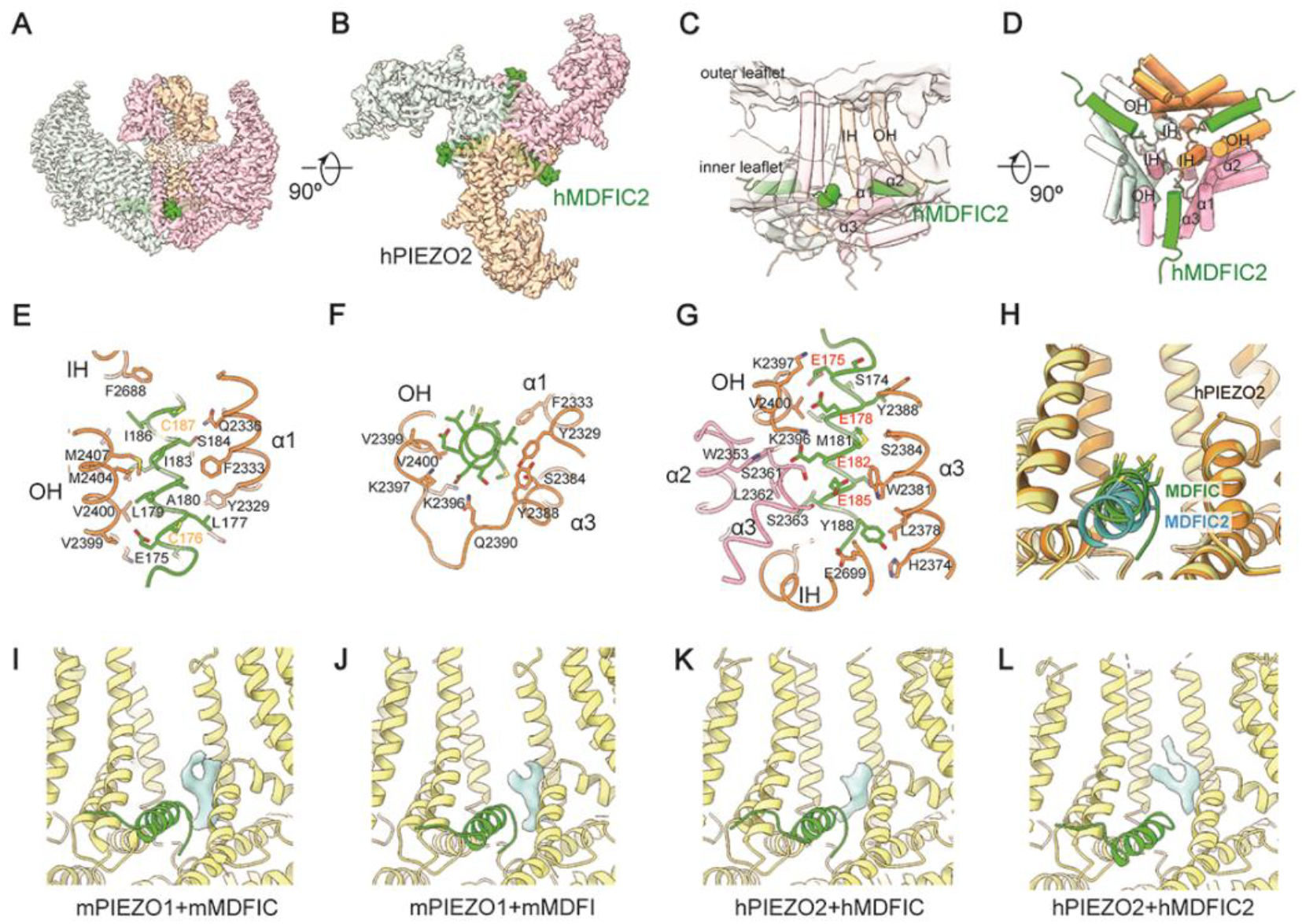
Structural characterization of an analogous unexplored auxiliary subunit binding site in human PIEZO2 for MyoD-family inhibitor proteins. (A-B) Cryo-EM density map of human PIEZO2:MDFIC2 complex. hMDFIC2 is coloured green and viewed from lateral (A) and cytoplasmic side (B). (C-D) The cysteine-rich C-terminal helix of hMDFIC2 lays parallel at the cytoplasmic side of the membrane, surrounded by the OH and anchor domain containing α1 to α3 (C) and extending deep into pore module of hPIEZO2 (D). (E-G) The interaction sites between hPIEZO2 and hMDFIC2, viewed from the membrane (E), lateral side (F), and cytoplasmic side (G). (H) MDFIC2 is positioned lower than MDFIC on PIEZO2. The model of hPIEZO2 (gold) in complex with MDFIC (green) is superimposed with the model of hPIEZO2 (yellow) in complex with MDFIC2 (cyan). The side chains of the cysteines in the C-terminal helix of MDFIC are shown as sticks. (I-L) Side views of the published structure of (I) mPIEZO1 with mMDFIC (PDB:8IMZ) and the new structures of all other MyoD-family inhibitor proteins (green) bound to PIEZOs (yellow) including (J) mPIEZO1 with mMDFI and (K) hMDFIC and (L) hMDFIC2 with hPIEZO2. The density of a trapped lipid between the MyoD-family inhibitor proteins and PIEZOs is shown in cyan.

Currently only one mouse PIEZO2 structure has been solved (45) showing a broadly similar architecture to mouse PIEZO1. Our complex of hPIEZO2:hMDFIC2 is the first structure of human PIEZO2, which has a consistently narrower pore configuration than that of human and mouse PIEZO1 (**Supplementary Fig. 6E**). We resolved the distal C-terminal helix of MDFIC2 which was situated at the bilayer interface (**Fig. 6C)** and intercalated into the PIEZO2 pore domain (**Fig. 6D)**, forming a network of interactions with the IH, OH, and α1–3 helices of the anchor domain (**Fig. 6E-G)**. When we compared the complexes of MDFIC and MDFIC2 bound to human PIEZO2 we noticed that MDFIC2 is positioned slightly lower than MDFIC on PIEZO2 (**Fig. 6H**) although this different positioning did not interrupt any of the important side chain interactions (**Fig. 6E-G)**.

When we compared all structures of MyoD-family inhibitor proteins with PIEZO1/2 we noticed there was an obvious lipid like density trapped between the MyoD-family inhibitor protein and the PIEZO1/2 pore shown in cyan. This lipid, which is absent in apo-PIEZO1/2 structures, would likely have to be pulled out prior to gating and may underlie the increased threshold of PIEZOs seen in the presence of all MyoD-family inhibitor proteins (**Fig. 6I-L**).

We also noticed additional density in the vicinity of two cysteines in the distal C-terminus of MDFIC2 that could not be explained by the protein alone (**Supplementary Fig. 6D**). So, we next investigated whether MDFIC2, like its relative MDFIC, underwent post-translational modification (PTM).

### MyoD-family inhibitor proteins regulate PIEZOs through a conserved mechanism

Previous work suggests that MyoD-family inhibitor proteins regulate PIEZOs through their C-terminal domain that undergoes palmitoylation on several key cysteine residues (25). This PTM is necessary for both the interaction with, and the regulation of, PIEZOs so we asked whether MDFIC2 also undergoes palmitoylation. Like MDFI and MDFIC, MDFIC2 migrated as two species on western blots (**Fig. 7A**). Palmitoylated proteins, despite the increase in size, tend to run further and hence look smaller on western blots. This was confirmed using an “unpalmitoylatable” MDFIC mutant where all potentially palmitoylated cysteines are mutated to alanine (MDFIC 15C-A) (**Fig. 7B**). This only runs as one band at the same size as the upper band in WT MDFIC. We biochemically confirmed palmitoylation of MDFIC2 using a PEG-maleimide mass-tagging assay. This involves using hydroxylamine (HAM) to cleave the palmitoyl chains and then reacting the remaining free-SH group with a pegylated maleimide so that the size of the protein increases for each palmitoylated cysteine. This experiment showed unequivocally that MDFIC2 undergoes palmitoylation on at least 2 cysteines, labelled as 1 palm and 2 palm (**Fig. 7C**). Comparison of the cryo-EM maps obtained for the hPIEZO2:hMDFIC and hPIEZO2:MDFIC2 structures revealed that less additional cysteine-associated density is observed on MDFIC2 compared to MDFIC; specifically, only Cys176 and Cys187 carried extra density for MDFIC2 (**Fig. 7D-E**). This may contribute to the lower positioning of MDFIC2 on PIEZO2 relative to MDFIC (**Fig. 6H**), possibly due to weaker interactions with the membrane. We then mutated these two cysteines to alanine and deleted the C-terminal 20 amino acids of MDFIC2 and both mutants failed to regulate stretch-evoked PIEZO1 currents (**Fig. 7F-H**). Therefore, akin to MDFIC/MDFI the sensory member of this family of proteins, MDFIC2, utilizes a palmitoylated, cysteine-rich C terminus to regulate PIEZO ion channels.

**Figure 7.**
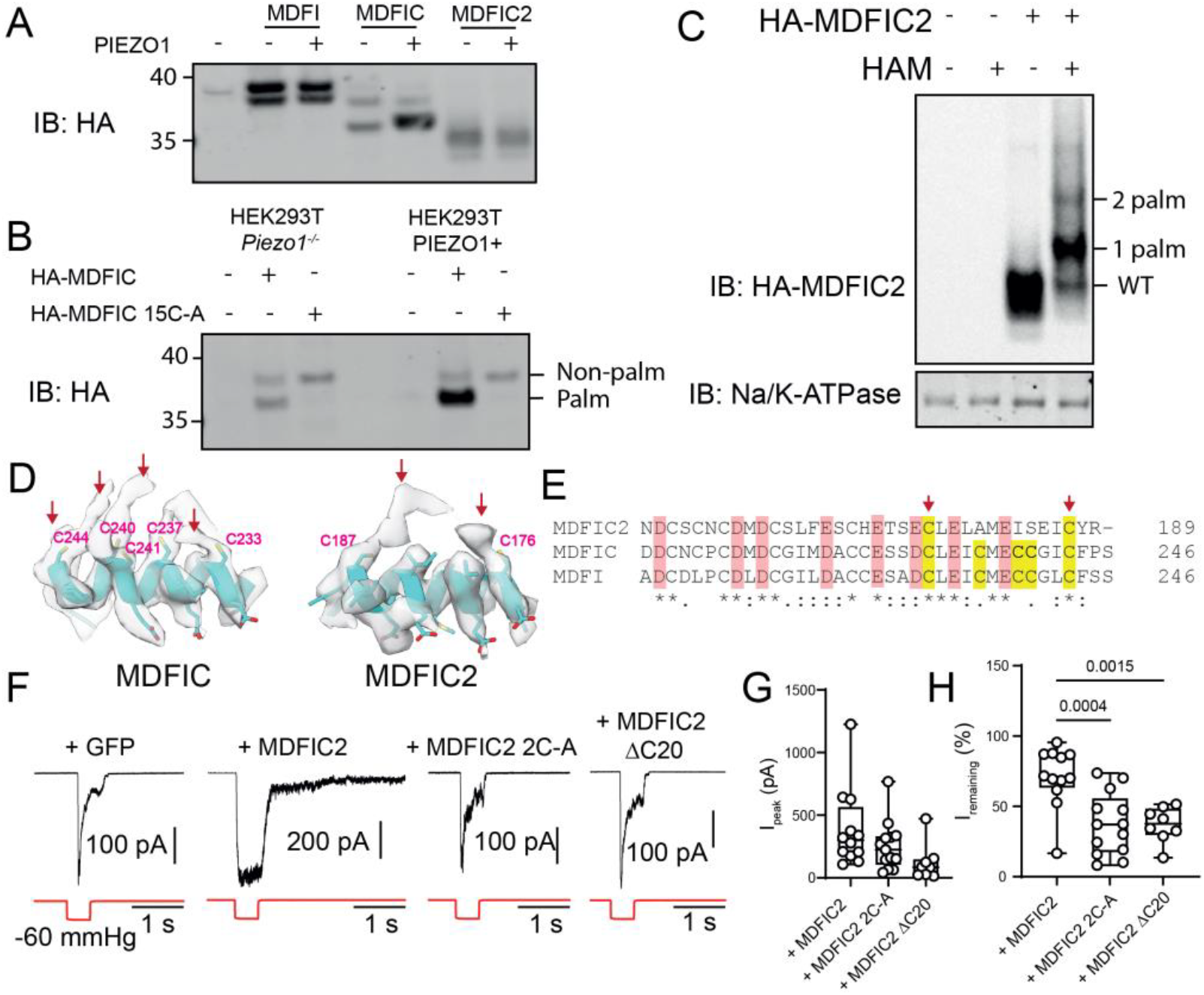
Cysteine palmitoylation in the C-terminus of MDFIC2 is essential for regulating PIEZOs. (A) Representative western blot of HA-tagged MDFI, MDFIC and MDFIC2 showing all three proteins run as two bands. (B) Comparison of WT MDFIC with an unpalmitoylatable mutant MDFIC (MDFIC15C-A) that only runs as a single band. (C) PEG-Maleimide mass tagging of MDFIC2 showing at least two cysteines undergo palmitoylation (HAM – Hydroxylamine; Palm - palmitoylation). (D) Comparison of C-terminal density of MDFIC and MDFIC2. (E) Sequence alignment of the distal C-terminus of MyoD-family inhibitor proteins highlighting the two cysteines with extra electron density in the MDFIC2 structure. (F) Representative cell-attached recording of human PIEZO1 alone compared to in the presence of WT MDFIC2, MDFIC2 2C-A(Cys176Ala/Cys187Ala) and MDFIC2 ΔC20 (deletion of the last 20 amino acids). (G-H) Quantification of replicate recordings of those shown in panel F for the peak current (G) and the current remaining at the end of the pressure pulse (H).

## Discussion

PIEZO channels are critical transducers of mechanical stimuli, converting physical forces into cellular signals across diverse tissues (18). In this study, we provide the first structural and functional characterization of an entire family of auxiliary subunits with distinct tissue-specific expression patterns. These subunits modulate key biophysical properties of PIEZO1 and PIEZO2 channels, including their inactivation profiles and mechanical activation thresholds, revealing a previously unrecognized layer of regulation that fine-tunes mechanosensory signalling across cell types.

In both sensory and non-sensory primary cell types, PIEZOs’ gating kinetics often diverge from those observed in heterologous expression systems. Most notably this manifests as slower inactivation kinetics (12, 13, 15, 18, 24). The slowly inactivating currents initially raise the possibility of the existence of other mechanosensitive ion channels with distinct kinetics. However, here we found instead of alternate pore-forming subunits, the slowly inactivating mechanically evoked currents from many cell lines and primary cells was carried by PIEZO1/2 channels complexed with MyoD-family inhibitor proteins.

The lack of RNA expression of *Mdfi* and *Mdfic* in sensory cell types led us to identify, and characterize a novel family member, MDFIC2, which is selectively and abundantly expressed in a subset of sensory cells. In DRG neurons these cells are marked by their strong expression of *Mrgprd* while being IB4 positive. These neurons are usually classified as high threshold mechanoreceptors and polymodal nociceptors (31, 46). Indeed, the biophysical phenotype of PIEZO1/2:MDFIC2 complexes, marked by a high mechanical activation threshold and sustained response due to loss of inactivation, is well-suited to the function of high-threshold mechanoreceptors (HTMRs). This is particularly relevant to the co-expression of MDFIC2 and PIEZO2 in *Mrgprd*+ DRG neurons, which exhibit slowly inactivating mechanically evoked currents at elevated thresholds that at least in part depend on PIEZO2 (6, 31). It strongly suggests that some of the slower adapting currents in DRGs and other sensory cells, including within the trigeminal ganglia, are complexes of PIEZO2 and MDFIC2.

Overexpression of MDFIC2 in the presence of native PIEZO2 in primary DRG neurons converts rapidly adapting currents to slowly adapting currents and suggests that the relative abundance of PIEZO2 compared with MDFIC2 will dictate the degree to which the inactivation is slowed. The effect on PIEZO2 inactivation in primary DRGs was much larger than that seen in PIEZO2 heterologously expressed in *Piezo1*^*-/-*^ N2A cells. This may suggest that other native proteins in the local environment collaborate with the PIEZO2:MDFIC2 complex to influence the final biophysical phenotype. We began to validate such a hypothesis by utilizing the PIEZO sensitizer STOML3 which lowered the gating threshold of PIEZO2:MDFIC2 thus increasing the proportion of slowly inactivating currents. Here it is important to note that multiple lines of evidence including patchseq data (28) suggest MDFIC2 (and indeed PIEZOs) cannot explain all slowly inactivating currents in DRG neurons. In Parpaite *et al*., 2021 many cells surveyed that had slowly inactivating current components lacked MDFIC2 and/or PIEZO expression (28). This is consistent with other entities acting as the pore forming subunits and contributing to slowly inactivating mechanosensitive currents.

What is the biological significance of MDFIC2 in regulating the kinetic properties of PIEZO channels? The inactivation of ion channels plays a critical role in precisely tuning ion flow. Modification of PIEZO1/2 channel inactivation has been widely linked to disease (47, 48). Therefore, the strong modulatory effect of MDFIC2 on PIEZO2 and their co-expression in functionally essential sensory neurons is likely to reshape PIEZO2-mediated mechanosensory processes. This requires future interrogation using animal models lacking *Mdfic2* expression. However the specific expression pattern in non-peptidergic DRG neurons (32) and the reduction of *Mdfic2* in models of pain (33, 34) would be consistent with a role in certain types of pain sensation.

Regulation of PIEZO2 by MDFIC2 occurs through a binding interface between adjacent subunits within the pore module in an analogous site to PIEZO1 (25). In fact, our suite of structures shows that the same site can be occupied by MDFI, MDFIC and MDFIC2, highlighting a shared mechanism of interaction through their palmitoylated C-terminus. Our cryo-EM structures represent the first glimpse of human PIEZO2 and identify a druggable interface with the potential for rational therapeutic design.

In summary, we report the identification and characterization of a novel PIEZO channel auxiliary subunit that is selectively and abundantly expressed in sensory cell types. We also present, to our knowledge, the first structural complex of human PIEZO2 with a binding partner. Given MyoD-family inhibitor proteins bind to not only PIEZO1/2 but also a broad range of transcription factors (40, 41, 49) we propose renaming this protein family as PTAPs (<u>P</u>iezo and <u>T</u>ranscription factor <u>A</u>uxiliary <u>P</u>roteins) to better reflect their molecular function. Our findings establish PTAPs as a unique family of pan-tissue auxiliary subunits that offers new insights into sensory physiology, pain mechanisms and beyond.

## Acknowledgements

We thank the Cryo-EM center at the Interdisciplinary Research Center on Biology and Chemistry, Shanghai Institute of Organic Chemistry for help with data collection. We thank the staff at the KGLMF at the University of New South Wales for processing the RNAScope samples.

## Author Contributions

Z.Z and M.L performed biochemical experiments. Z.Z, C.D.C, Y.G & D.C. performed siRNA knockdown and electrophysiology. J.C, & E.W. analysed scRNAseq data. D.C carried out imaging. Z.Z, M.L., R.L, J.V.L & S.F.O. generated reagents and isolated primary cells. F.D, X.M, H.Z, and Y.Z performed cryo-EM structural studies. Y.Z and C.C supervised the project. Y.Z and C.C conceived of the project. All authors wrote and approved the manuscript.

## Data availability

The composite map of the mPIEZO1-MDFI, hPIEZO2-MDFIC, hPIEZO2-MDFIC2 complexes have been deposited in the Electron Microscopy Data Bank under the accession codes EMD-64998, EMD-65006 and EMD-65005 respectively. The atomic coordinates have been deposited in the Protein Data Bank under the accession codes 9VED, 9VEF, and 9VEE. The composite map may contain artificial features near the boundaries of the masks. The consensus maps, masked refined cap maps, and masked refined TMD maps have been deposited under the accession codes EMD-64999, EMD-65000, EMD-65001, EMD-65002, EMD-65003, and EMD-65004. All data are available in the main text or the supplementary materials.

