## Supporting Information for "MDFIC2 is a sensory neuron-specific PIEZO channel auxiliary subunit"

*^1^Victor Chang Cardiac Research Institute, Sydney, NSW, 2010, Australia. ^2^ School of Clinical Medicine, Faculty of Medicine and Health, University of New South Wales, Sydney, NSW 2052, Australia. ^3^Interdisciplinary Research Center on Biology and Chemistry, Shanghai Institute of Organic Chemistry, Chinese Academy of Sciences, China. ^4^School of Biotechnology and Biomolecular Sciences, Faculty of Medicine and Health, University of New South Wales, Sydney, NSW 2052, Australia. ^5^Shanghai Key Laboratory of Aging Studies, Shanghai, 201210, China.* *^6^State Key Laboratory of Chemical Biology, Shanghai Institute of Organic Chemistry, Chinese Academy of Sciences, Shanghai, 200032, China. ^7^School of Biomedical Sciences, Faculty of Medicine & Health, UNSW Sydney, Kensington, NSW, 2052, Australia*

*These authors contributed equally

#Corresponding authors:

**Materials and Methods**

**Antibodies and plasmid constructs**

Sources and details of all antibodies used in this article are summarized in Table S2. Sources and details of all plasmid constructions in this article are summarized in Table S3. To generate plasmids expressing MDFIC2 with an N-terminal HA tag the coding sequences were directly synthesized as gBlocks (IDT). The DNA fragments were then digested and annealed into plasmid backbones. To generate mutations or truncated versions of MDFIC2 site-directed mutagenesis was undertaken using a custom protocol with the high-fidelity polymerase PfuUltra.

**Cell culture and transfection**

*Piezo1^−/−^* HEK293T cells were a gift from Dr. Ardem Patapoutian (The Scripps Research Institute, La Jolla, CA, United States); Neuro2A (N2A) cells were a gift from Dr Kate Poole (UNSW, Sydney, Australia) and *Piezo1^−/−^* N2A cells were a kind gift from Dr Gary Lewin (MDC, Berlin). HL-1 cells were a kind gift from Dr Richard Harvey (VCCRI, Sydney). 3T3-L1 cells were a kind gift from Dr Renjing Liu (VCCRI, Sydney). MCC13 cells were purchased from CellBank Australia (Cat# 10092302). HOG cells were purchased from Sigma (SKU# SCC163). HeLa cells were a kind gift from Dr Boris Martinac (VCCRI, Sydney). HAEC were purchased from Lonza (Cat# CC-2535). Cell lines were not authenticated and aside from HeLa were not listed in the Database of commonly misidentified cell lines maintained by ICLAC (http://iclac.org) and NCBI Biosample (<http://www.ncbi.nlm.nih.gov/biosample>). N2A, HEK293T, 3T3-L1, HeLa, and HOG cells were maintained in DMEM supplied with 10% FBS and Penicillin-Streptomycin (Gibco, 15140122). MCC13 cells are maintained in RPMI supplied with 15% FBS and Penicillin-Streptomycin (Gibco, 15140122). HL-1 cells are maintained in Claycomb media. Cells were transfected with polyethylenimine (PEI) or Lipofectamine 3000 transfection reagent (Thermo Fisher Scientific).

**Mouse lung endothelial cell** **isolation**

Mouse lung endothelial cells were isolated based on protocol by Wang et al (1). Briefly, lungs were dissected from 3–4-week-old mice and digested in collagenase type I (2 mg/ml) solution for 1 h at 37 °C. Digested tissue was filtered through 70 μm cell strainer and endothelial cells purified using Dynabeads (Invitrogen) incubated with CD31 antibody (BD Pharmingen). Isolated cells were cultured in EGM-2 medium with media changed every 2-3 days. Cells were transfected with siRNA in RPMI with 1% FBS without antibiotics.

**Immunoprecipitation**

To immunoprecipitate MDFI, MDFIC and MDFIC2 and co-immumoprecipitate PIEZO1-HaloTag, PIEZO2-mCherry or FLAG-GATA2, plasmids were transfected into *Piezo1^−/−^* HEK293T cells and the cells were lysed in CHAPS lysis buffer (1% w/v CHAPS, 0.6% w/v soy PC, 140 mM NaCl, 1 mM EDTA, 25 mM NaPIPES) supplemented with 2 mM 1,4-Dithiothreitol (DTT; Sigma) and 1× protease-inhibitor cocktail (Roche) 48 h post transfection. Lysates were centrifuged at 13,200 rpm for 20 min at 4 °C; the expression of each protein of interest was confirmed by immunoblotting 5% of the collected supernatants, and the remaining supernatants were incubated with indicated antibodies and Protein G Dynabeads (Thermo Fisher Scientific), or HA-Trap magnetic agarose (ChromoTek) at 4 °C overnight. The recovered beads were washed three times with CHAPS wash buffer (25 mM NaPIPES, 140 mM NaCl, 0.6% w/v CHAPS, 0.14% PC) supplemented with 2 mM 1,4-Dithiothreitol (Sigma) and 1× protease-inhibitor cocktail (Roche), and heated in 2x sodium dodecyl sulphate (SDS)-PAGE loading buffer containing 1 M urea and 10 mM tris(2-carboxyethyl) phosphine (TCEP) at 55°C for 5 min. Both the total cell lysate (input) and the eluted proteins were subjected to western blot analysis.

**Western blotting**

Cells were solubilized in a modified RIPA buffer [Tris buffer 10 mM, ethylenediaminetetraacetic acid (EDTA) 1 mM, NaCl 140 mM, in (% w/v): Sodium deoxycholate 0.1, SDS 0.1, Triton X-100 1.0, pH 7.2] supplemented with 1× EDTA-free protease inhibitor cocktail tablets (Sigma-Aldrich) and 2 mM DTT for 10 min on a rotating wheel at 4 °C unless otherwise specified. Cell lysates were cleared by centrifugation at 13,000 ×g at 4 °C for 10 min. Supernatant was mixed with SDS loading buffer and subjected to 3-8% Tris-Acetate or 4-12% Bis-Tris SDS PAGE (Invitrogen). Proteins were transferred to nitrocellulose membranes (Bio-Rad) and the membrane was blocked with 4% non-fat milk in TBST for 1 h. The membrane was then blotted with primary antibody overnight at 4°C and subsequently with a secondary antibody for 1 h (All antibodies can be found in Table S2). Protein bands were detected with Li-Cor Odyssey Infrared Imaging System 9120 (Li-Cor) or Chemidoc MP imaging system (Bio-Rad).

**Native gel electrophoresis and western blot**

Transfected HEK293T cells were lysed with a low salt lysis buffer (1% w/v CHAPS, 0.6% w/v soy PC, 50 mM NaCl, 1 mM EDTA, 25 mM NaPIPES) supplemented with 2 mM 1,4-Dithiothreitol (DTT; Sigma) and 1× protease-inhibitor cocktail (Roche). Lysates were centrifuged at 13,200 rpm for 10 min at 4°C. Native gel electrophoresis and western blot were conducted according to manufacturer’s instructions. In brief, supernatant was mixed with NativePage sample buffer (Thermo Fisher Scientific) and G-250 sample additive (Thermo Fisher Scientific) then loaded to a 4-16% NativePage (Thermo Fisher Scientific). After gel electrophoresis, marker lane was cut and stained with Coomassie blue reagent, and protein samples were transferred to a nitrocellulose membrane. Proteins on the membrane were fixed with 8% acetic acid for 10 min at room temperature. The membrane was then subjected to immunoblotting following the protocol as suggested above except for an HRP-conjugated secondary antibody was used. The membrane was then briefly incubated with Pierce™ ECL Western Blotting Substrate (Thermo Fisher Scientific) and protein bands were detected with a ChemiDoc MP Imaging System (Bio-Rad).

**Protein labelling for confocal microscopy**

Cells were dissociated with TrypLe express enzyme (12604013, Gibco) and plated on CellCarrier Ultra microplate (PerkinElmer) coated with fibronectin 3.3 µg/ml (Sigma) in 0.2% Gelatin in PBS overnight. The cells were fixed with 4% paraformaldehyde in PBS for 30 minutes at room temperature. Samples were then washed with PBS then quenched with 0.1 M Glycine in Tris-buffered saline (TBS) for 10 min. The cells were then permeabilized with 0.1% Triton-X100 in TBS for 5 min and blocked with 5% bovine serum albumin (BSA) for 1 h at room temperature.

To label the different proteins, the cells were incubated with primary antibody (refer to Table S2) diluted in 3% BSA for 1 h then an appropriate secondary antibody for 1 h, at room temperature. Nuclei were labelled with DAPI and, when necessary, cell membranes were labelled with 2 µg/ml of wheat germ agglutinin–AF647 (Biotium) for 20 min at room temperature. Finally, the samples were imaged using a confocal microscope and a 60x oil objective (LSM900, ZEISS).

**siRNA mediated knockdown**

To account for the slow turnover of PIEZO proteins and maximize knockdown efficiency two treatments with dsiRNA (IDT) were performed. 10 nM of negative control dsiRNA, or combinations of PIEZO1/2 or MDFIC/MDFI dsiRNAs as indicated in the text, were transfected using RNAiMAX (Invitrogen) according to the manufacturer’s instructions. Targeting sequences of dsiRNA used are: *mPiezo1*: auccgugccaaacaggag; *mPiezo2*: cgagaggaugagccagu; *mMDFIC*: guuuaucuauuggagguu; *mMDFI*: uucgauucauuacuguaa; *hPiezo1*: accaagaaguacaaucau; *hPiezo2*: ucgaaagaauaucgcuaa; *hMDFIC*: cugagaaagauauaacuc; *hMDFI*: gaguuccugacgcugug. After 48 h cells were lifted and transfected again with the same amount of transfection reagents and dsiRNA. Cells were then cultured for another 48 h before being seeded for electrophysiological recordings.

**Electrophysiology**

Cells were plated on gelatin coated glass coverslips for cell-attached patch clamp analysis. The extracellular solution was: 90 mM K^+^Aspartate, 50 mM KCl, 2 mM MgCl_2_ and 10 mM HEPES (pH 7.2) adjusted with KOH in order to zero the membrane potential. The pipette solution contained either: 90 mM CsCl, 40 mM CsF, 1 mM EGTA with 10 mM HEPES (pH 7.2) adjusted with CsOH or for TREK-1 recordings only 140 mM KCl, 1 mM CaCl_2_, 1 mM MgCl_2_ and 10 mM HEPES adjusted to pH 7.2 using KOH. Negative pressure was applied to patch pipettes using a High-Speed Pressure Clamp-1 (ALA Scientific Instruments, Farmingdale, NY, United States) and recorded in millimetres of mercury (mmHg) using a piezoelectric pressure transducer (WPI, Sarasota, FL, United States). Each pressure pulse sweep was separated by a 50 s rest interval to maximize the chance for all channels to close between sweeps. Borosilicate glass pipettes (Sigma-Aldrich) were pulled with a vertical pipette puller (PP-83, Narashige, Tokyo, Japan) to produce electrodes with a resistance of 2.5-3.5 MΩ. The currents for PIEZO1 or PIEZO2 were amplified using an AxoPatch 200B amplifier (Molecular Devices, LLC), and Data were sampled at a rate of 10 kHz with 1 kHz filtration and analyzed using pCLAMP10 software (Molecular Devices, LLC).

For whole-cell recordings transfected cells were plated onto poly-L-lysine (Sigma) and fibronectin (150 nM, Sigma)-coated coverslips prior to electrophysiological experiments. Flatter cells like endothelial cells we plated on ~5 kPa PDMS layered coverslips coated with fibronectin (150 nM) and laminin (10 μg/ml, BioLamina). For DRGs, glass coverslips were coated with fibronectin (150 nM) and laminin (10 μg/ml). Cells were indented using a fire-polished glass pipette with a tip diameter of 2-3 μm controlled by a piezo-electric driver (E625 servo Controller/Amplifier; Physik Instrumente) mounted on an MP285 manipulator (Sutter instruments) using a custom 3D printed bracket. The internal solution contained in mM: 133 CsCl, 10 HEPES, 1 EGTA, 2 MgCl_2_, 4 MgATP, 0.4 Na_2_GTP, pH 7.3 with CsOH. The bath solution was 130 NaCl, 3 KCl, 1 MgCl_2_, 10 HEPES, 2.5 CaCl_2_, 10 glucose (pH 7.3 with NaOH). Electrophysiological recordings were acquired at 10 kHz using a Digidata 1440A digitizer (Molecular Devices, LLC), an Axopatch 200B amplifier (Molecular Devices, LLC) filtering (4-pole Bessel filter) at 5 kHz using the patch software pCLAMP10 (Molecular Devices, LLC). Clampfit 11.1 (Molecular Devices, LLC) was used to analyze the data including for all bi- and mono-exponential fits of inactivation time constants. All data were acquired from at least three independent cell transfections. All patch-clamp recordings were performed at room temperature >48 h after transfection.

The Boltzmann plots were obtained by fitting PIEZO2 channel open probability *P*_o_∼*I*/*I*_max_ versus negative pressure using the expression *P*_o_/(1–*P*_o_)=exp [*α*(*P*–*P*_50_)], where *P* is the negative pressure [mmHg], *P*_50_ is the pressure at which *P*_o_=0.5, and *α* [mmHg^−1^] is the slope of the plot ln [*P*_o_/(1–*P*_o_)=[*α*(*P*–*P*_1/2_)] which reflects channel mechanosensitivity.

**Quantitative PCR**

Total RNA was extracted using TRI Reagent (Sigma), and cDNA was synthesized using High-Capacity cDNA Reverse Transcription Kit (Invitrogen). Quantitative PCR (qPCR) amplifications of various genes were performed using SYBR Green (Takara) and a CFX384 Touch Real-Time PCR Detection System (Bio-Rad); GAPDH/HPRT were used as normalization controls. After initial denaturation (30 s, 95 °C), amplification was performed using 40 cycles of 5 s at 95 °C and 30 s at 60 °C.

The sequences of the primers used for qPCR were the following: *mMDFIC2* (F: ACACCTCTGGCTTCCTCCTT, R: TTTCCTGGGCTGGTCCATCTG); *mPiezo2* (F: CCATCTTTGACCTCGGCTGT; R: CCATCTTTGACCTCGGCTGT); *mHPRT1* (F: GCTTGCTGGTGAAAAGGACCTC; R: CCTGAAGTACTCATTATAGTCAAGGGC); *mGAPDH* (F: TATGTCGTGGAGTCTACTGG, R: AGTGATGGCATGGACTGTGG).

**Isolation of mouse DRGs, TGs and JNGs**

All experimental procedures were approved by the Garvan Institute/St. Vincent’s Hospital Animal Experimentation Ethics Committee. All experiments were conducted in accordance with the Australian code for the care and use of animals for scientific purposes. Mice were housed on a 12-h light/ 12-h dark cycle and provided with normal chow diet and water *ad libitum*.

The protocol for mouse DRG neurons isolation is adapted from previous work (2). Briefly, mice aged 6-8 weeks were anesthetized with isoflurane, followed by cervical dislocation. The spin was separated and cut open in cold HBSS, and 30 to 40 DRG nodes were isolated from one animal. The DRG nodes were briefly washed with 10 ml of cold DMEM with 10% FBS, then digested with Liberase TL, at a concentration of 125 μg in 1 ml of serum-free DMEM for 1 h in a 37 °C incubator. The tissue was gently agitated every 10 min. After this, 125 μg of Liberase TM was added to the media, and digestion continued in the incubator for 10 min.

Once the digestion was finished, 2 ml of DMEM with 10% FBS was added to the tissue, and the tissue was homogenized by pipetting up and down around 15-25 times. The mixture was passed through a 100 μm cell strainer and spun down at 250 x g for 5 min to pellet the cells. For one animal, the pellet was resuspended in 200 μL of DRG culture media (DMEM, 10% FBS, 50 ng/ml NGF (Gibco), 25 ng/ml GDNF (Gibco), penicillin/streptomycin) and 100 μL of cell resuspension was carefully added to a well in a 12-well plate, pre-coated with fibronectin (150 nM) and laminin (10 μg/ml). After one hour of cell seeding, 1 ml of DRG culture media was added to the well. DRG neurons were maintained for 1 to 3 days for the following experiments.

Mouse TGs and JNGs were dissected following previously described protocols (3) (4). For RNAScope, isolated DRGs, TGs and JNGs were immersed fixed in 4% paraformaldehyde overnight then infiltrated in 40% sucrose, embedded in OCT and frozen in liquid nitrogen for subsequent RNA detection and fluorescence labelling.

**RNAScope**

Frozen sections of 8 µm were taken from the ganglia and processed for RNA detection using the RNAScope™ Multiplex Fluorescent Reagents Kit (PN 323100, ACD) following the manufacturer’s protease-based protocol. Nuclei were labelled with DAPI and sections were mounted in ProLong Gold Antifade mounting media (P36934, Invitrogen) before being imaged using a confocal microscope and a 40x objective (LSM900, ZEISS). The RNAScope probes used were: Mm-Md-Fic2-O1-C2 (1779041-C2) and Mm-Piezo2-E43-E45-C3 (439971-C3), combined with Opal dye 570 (FP1488001KT) and Opal dye 690 (FP1497001KT) respectively, at 1:1000. Consecutive sections were labelled with Griffonia Simplicifolia Lectin I (GSL I) Isolectin B (IB4) conjugated to Alexa Fluor568 (I21412, Invitrogen) at 1:200.

**Mouse DRG electroporation**

After culturing for one or two days, the cells were gently washed with PBS once to remove debris. The cells were then lifted with trypsin and counted. Cells (6x10^4^) were resuspended in 10 μL of buffer R, mixed with 1000 ng of plasmid, and subjected to electroporation with the Neon electroporation system (1200 V, 200 ms, 2 pulses). Cells were then seeded onto fibronectin/laminin-coated glass coverslips with recovery media (alpha-MEM (Gibco), 1% FBS, 1% N2 supplement) and 50 ng/ml NGF, 25 ng/ml GDNF, and penicillin/streptomycin were added after 2 h of seeding. The transfected DRGs were used within 24 to 72 h after electroporation.

**Protein expression and purification**

Mouse PIEZO1 and human PIEZO2 genes were cloned individually into pEG Bacmam vector (5) with a HRV-3C cleavable GFP tag at the C-terminus. Mouse MDFI, human MDFIC, and human MDFIC2 genes were individually cloned into pEG Bacmam expression vector with a N-terminal Flag tag. Plasmids were transformed into DH10bac bacterial strain (Weidibio) to generate bacmids. Baculovirus was produced by transfecting bacmids into *Spodoptera Frugiperda 9* (Sf9) cells by Fugene HD (Promega) in ESF921 medium (Expression Systems). HEK293S GnTI^-^ suspension cells were cultured in SMM 293-TII Expression Medium (Sino Biological) or FreeStyle^TM^ 293 Expression Medium (Gibco) in a Herocell C1 orbital shaker incubator (Radobio) set at 37 °C with 8% CO2. Cells at a density of 2~3 x 10^6^ cells/ml were transfected with Piezo virus and corresponding MDFIC virus at a ratio of 1:1. After 16 h, 10 mM sodium butyrate was added to enhance protein expression and the culture temperature was adjusted to 30 °C. After another 48 h, transfected cells were harvested by centrifugation at 3,250 x g, 4 °C for 20 min.

For the mPIEZO1-mMDFI complex and hPIEZO2-hMDFIC complex, cells were lysed in lysis buffer containing 30 mM Tris-HCl, pH 7.9, 300 mM NaCl, 1 mM dithiothreitol, 1 µg/mL pepstatin, 1 µg/mL aprotinin, 1 µg/mL leupeptin, 20 µg/mL trypsin inhibitor, 2 µg/mL DnaseI, 2.5 mM MgCl_2_, 0.5 mM CaCl_2_, 3 mM PMSF, and 1% C_12_E_10_ at 4 °C for 2 h. Cell debris was then removed by centrifugation at 30,000 x g, 4 °C for 40 min. The supernatant was incubated with anti-Flag affinity resin at 4 °C for 2 h. Resin was collected by centrifugation at 300 x g, 4 °C and extensively washed with wash buffer containing 30 mM Tris-HCl, pH 7.9, 150 mM NaCl, and 0.05% GDN. Proteins were eluted with the same buffer supplemented with 0.25 mg/mL Flag peptide. Eluted proteins were concentrated using Ultra-15 100-kDa cutoff centrifugal filter (MilliporeSigma) and loaded onto a Superose 6 increase 10/300 GL column (Cytiva) in AKTA (Sepure) equilibrated in 30 mM Tris-HCl, pH 7.9, 150 mM NaCl, 0.02 % GDN. The peak fractions were pooled and immediately used for Cryo-EM sample preparation.

For the hPIEZO2-hMDFIC2 complex, the purification process was the same as above, except that the eluted proteins were directly concentrated for Cryo-EM sample preparation without further gel filtration due to the lower yield.

**Cryo-EM sample preparation and data collection**

Aliquots of 4 µl fresh protein were applied to glow-discharged (Harrick Plasma) Au grids (Quantifoil, R2/1, 200 mesh) in the chamber of a Vitrobot (Mark IV, Thermo Fisher Scientific) set at 100% humidity and 4 °C. After waiting for 5 s, grids were blotted for 2 s with a blot force of -2 and plunged into pre-cooled liquid ethane.

Cryo-EM data were collected on a 300 kV Titan Krios G4 microscope (Thermo Fisher Scientific) equipped with Biocontinuum K3 Direct Electron Detector with GIF energy filter (Gatan). Grids were screened, and movies were automatically acquired using EPU software. The movie stacks were collected in super-resolution mode at a nominal magnification of ×81,000 with a pixel size of 0.5275 Å and a defocus range of −1.2 to −2.2 μm. The dose rate was 25 e^−^ /pixel/s, and exposures of 2.5 s were dose-fractionated into 40 frames.

**Imaging processing**

For all samples, the movie stacks were first imported to RELION-3.1 (6) and motion corrected with MotionCor2 (7). The CTF parameters were determined with CTFFIND4 (8). Particles were automatically picked with Gautomatch (https://github.com/JackZhang-Lab/Gautmatch).

For the hPIEZO2-hMDFIC2 complex, extracted particles were initially imported into CryoSPARC (9). A small subset was used to generate initial references via ab initio reconstruction with C3 symmetry. Using these references, all extracted particles were subjected to heterogeneous refinement with C3 symmetry to remove junk particles. Particles from the well-defined classes were then imported into RELION-3.1 for Bayesian polishing. The polished particles were re-imported into CryoSPARC and processed through another round of heterogeneous refinement. The EM map from the well-defined class was selected as the initial reference for an additional round of heterogeneous refinement to distinguish different conformational or compositional states of the complex. Particles from the class with well-defined hMDFIC2 density were selected for non-uniform refinement with C3 symmetry. To improve the map quality of the cap and transmembrane domains, focused 3D classifications were performed using masks targeting these regions. Particles from classes with clear structural features were selected and subjected to local refinement. The two resulting locally refined maps were then combined to generate the final composite map of the complex. Due to the low occupancy of hMDFIC2, C3 symmetry expansion followed by hMDFIC2-masked 3D classification was conducted to assess the proportion of particles containing hMDFIC2 (Supplementary Figure. 6).

For the hPIEZO2-hMDFIC complex, extracted particles were first subjected to 2D classification in RELION-3.1. Particles from classes exhibiting clear hPIEZO2 features were selected and imported into CryoSPARC. A subsequent heterogenous refinement with C3 symmetry was performed using references generated from a small subset of particles. The class showing clear structural features was selected and subjected to three rounds of ab initio reconstruction with C3 symmetry to remove junk particles and enhance the density of hMDFIC. The selected particles were then imported back to RELION-3.1 for Bayesian polishing. The polished particles were re-imported into CryoSPARC and subjected to non-uniform refinement with C3 symmetry. Local refinements were performed with cap and transmembrane domain masks to further improve the map quality of these regions (Supplementary Figure. 7).

For the mPIEZO1-mMDFI complex, particles were initially processed by 2D classification in RELION 3.1 to discard obvious junk particles. Selected particles with recognizable secondary structure features were imported into CryoSPARC for ab initio reconstruction. The class displaying clear mPIEZO1-mMDFI features was then selected for non-uniform refinement (Supplementary Figure. 7).

**Model building**

For the hPIEZO2-hMDFIC complex and hPIEZO2-hMDFIC2 complex, AlphaFold2-predicted structures were used as initial models (10). These models were fitted into the Cryo-EM maps using UCSF Chimera (11). Three symmetric extra densities located near the pore module of hPIEZO2 were assigned as the C-terminal helices of hMDFIC or hMDFIC2. For the mPIEZO1-mMDFI complex, the previously reported structure of mPIEZO1-mMDFIC (PDB: 8IMZ) was used as starting model, with the mMDFIC region replaced by AlphaFold2-predicted C-terminal helix of mMDFI. All atomic models were subsequently refined using real-space refinement in PHENIX(12) and manually adjustment in Coot(13).

**Mass tagging pegylated maleimide**

For mass tagging with pegylated maleimide cell samples were incubated with a modified RIPA buffer [Tris buffer 10 mM, ethylenediaminetetraacetic acid (EDTA) 1 mM, NaCl 140 mM, in (% w/v): Sodium deoxycholate 0.1, SDS 0.1, Triton X-100 1.0, pH 7.2] supplemented with 1× EDTA-free protease inhibitor cocktail tablets (Sigma-Aldrich), 1 mM (phenylmethylsulfonyl fluoride) PMSF, 2 mM DTT, and 40 mM N-ethylmaleimide (NEM) overnight at 4°C. Protein were then precipitated by methanol-chloroform-H_2_O precipitation (4:1.5:3) with sequential addition of methanol (400 µL), chloroform (150 µL), and distilled H_2_O (300 µL) (all prechilled on ice). The reactions were then mixed by inversion and centrifuged (Centrifuge 5417R, Eppendorf) at 20,000 × g for 5 min at 4 °C. To pellet the precipitated proteins, the aqueous layer was removed, 1 mL of prechilled MeOH was added, the Eppendorf tubes were inverted several times and centrifuged at 20,000 × g for 3 min at 4 °C. The supernatant was then decanted, and the protein pellets were dried with an N_2_ stream. To completely remove the NEM, the protein pellets were resuspended with 4% SDS and re-precipitated three times. For hydroxylamine (HAM) cleavage and mPEG-maleimide alkylation, the protein pellet was resuspended in 60 µL binding buffer A containing 4% SDS, 4 mM EDTA and 30 µL resuspended protein was treated with 90 µL of 1 M neutralized HAM to obtain a final concentration of 0.75 M HAM. The other 30 µL resuspended protein was diluted in 90 µL binding buffer A containing 250 mM NaCl as control samples. Samples were incubated at room temperature for 1 h with nutation. The samples were then subjected to methanol-chloroform-H_2_O precipitation, as described above, and resuspended in 30 µL binding buffer A containing 4% SDS, 4 mM EDTA then treated with 90 µL binding buffer A containing 1.33 mM 10 kDa mPEG-Mal (Sigma) for a final concentration of 1 mM mPEG-Mal. Samples were incubated for 1.5 h at room temperature with nutation before a final methanol-chloroform-H_2_O precipitation. Dried protein pellets were resuspended in 50 µL 1× SDS loading buffer and then heated for 5 min at 95 °C. Typically, 25 µL of the sample was subjected to western blot analysis. All acyl biotin exchange and mass tagging were carried out with material from at least three independent transfections.

**Single-cell RNA-seq analysis**

**Processed single-cell RNA data from Ziesel et al., 2018 was download from** [**http://mousebrain.org/**](http://mousebrain.org/) **(l2_neurons_drg.loom), and data from Usoskin et al., 2015 was obtained from** [**http://linnarssonlab.org/drg/**](http://linnarssonlab.org/drg/)**. Both datasets were processed using the same parameters following log-normalization of counts using a scaling factor of 10,000 using Seurat (14). The top 2,000 highly variable features were used for principal component analysis followed by construction of UMAPS using the first 10 principal components. Cells clusters were identified using FindClusters with a clustering resolution of 0.5. For the Usoskin et al., 2015 data cell types were annotated using the provided meta-data after filtering of empty cells, doublets and outlier cells. Cell type annotation for the Ziesel et al., 2018 data was carried out based on the expression of maker genes identified using FindAllMarkers across clusters.**

**Statistical analysis**

Statistical analyses were performed, and significance was computed using GraphPad Prism 9. The specific test used is stated in the respective figure legend. All exact p-values <0.05 are shown, for values greater than 0.05 they are denoted by n.s (non-significant). Student’s t-test was used when comparison was made between two groups. One-way ANOVA with multiple comparison (Tukey’s) test was used when comparing three groups. All data are displayed as means ± SEM or maximum to minimum box and whiskers plots*.*

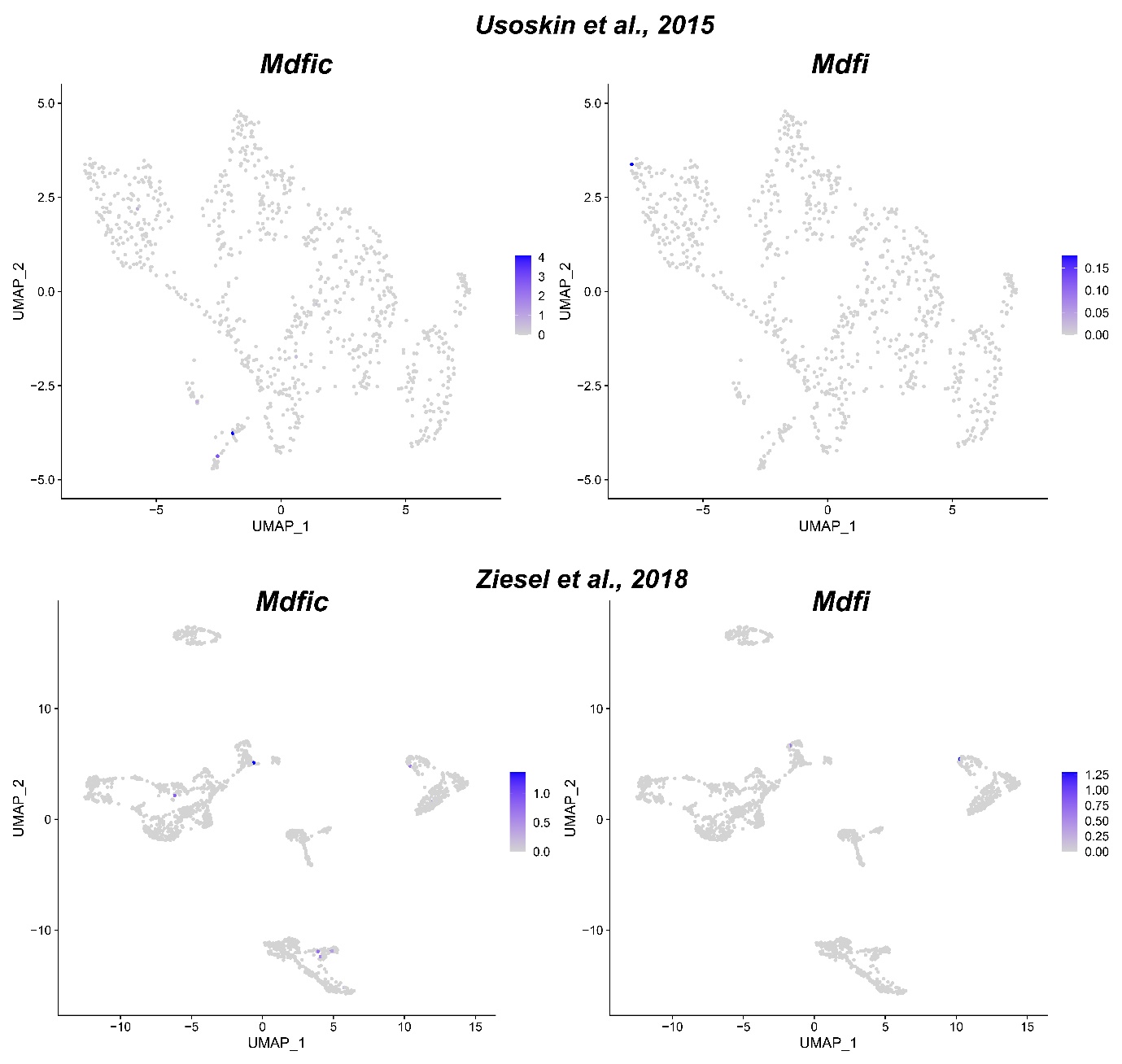

***Supplementary Figure 1. Mdfic and Mdfi are not expressed in any DRG subpopulations at noticeable levels.*** *Log-normalized expression data is shown for Mdfic and Mdfi captured from Ziesel et al., 2018.*

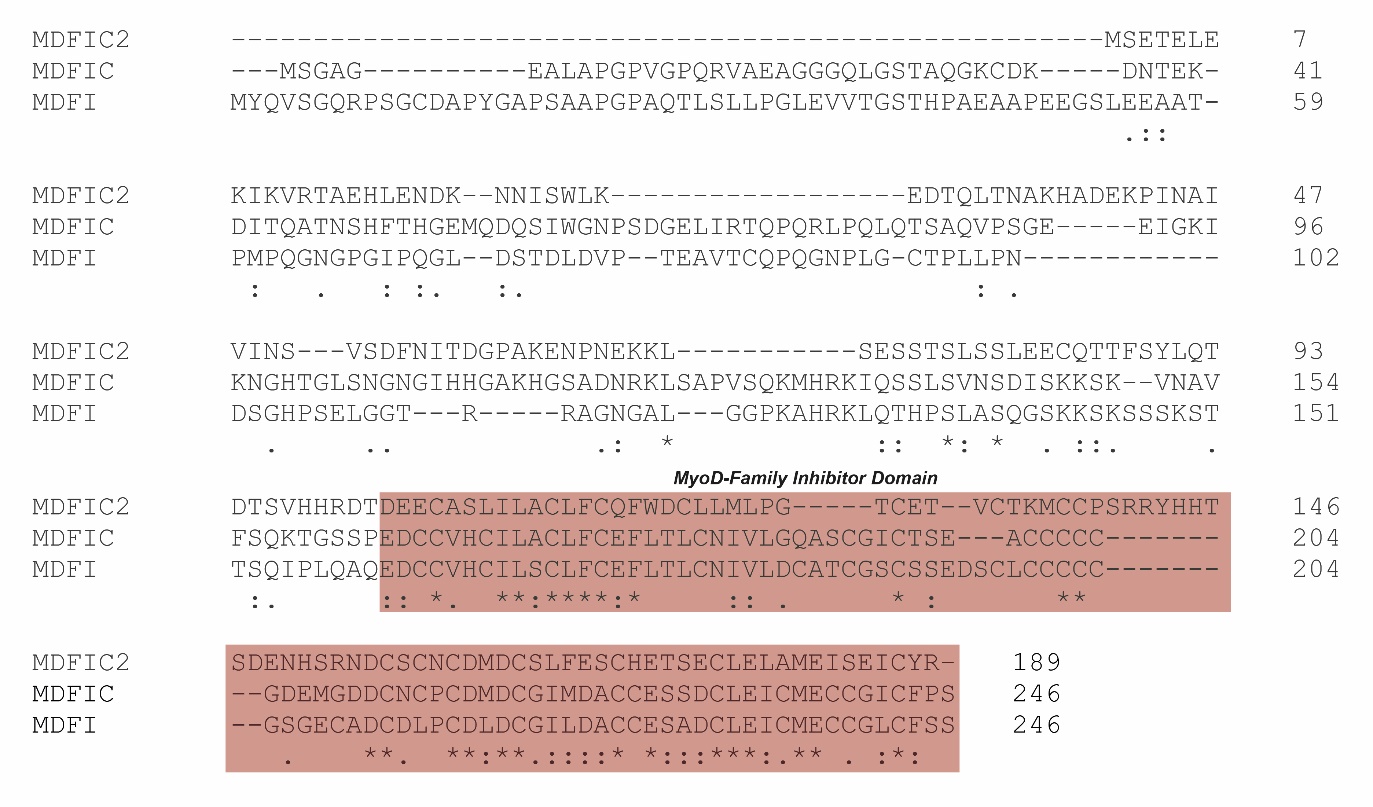

***Supplementary Figure 2. Sequence alignment of human MDFIC2 compared to human MDFIC and MDFI.*** *The “MyoD-Family Inhibitor Domain” is highlighted in red.*

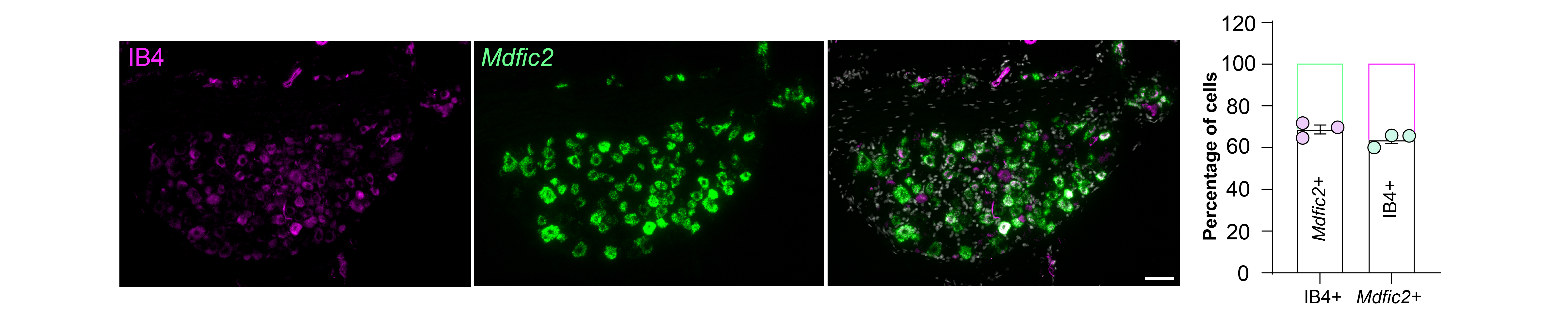

***Supplementary Figure 3. IB4 and Mdfic2 co-localize in DRGs.*** *RNAScope and fluorescence labelling data showing IB4 positive and MDFIC2 positive cells co-localize in a subset of DRG neurons (Scale bar – 50 μm). 69% of n=186 IB4 positive cells are MDFIC2+; 64% of n=199 MDFIC2 positive cells are IB4 positive. Statistics are from 3 individual DRGs. White represents nuclei.*

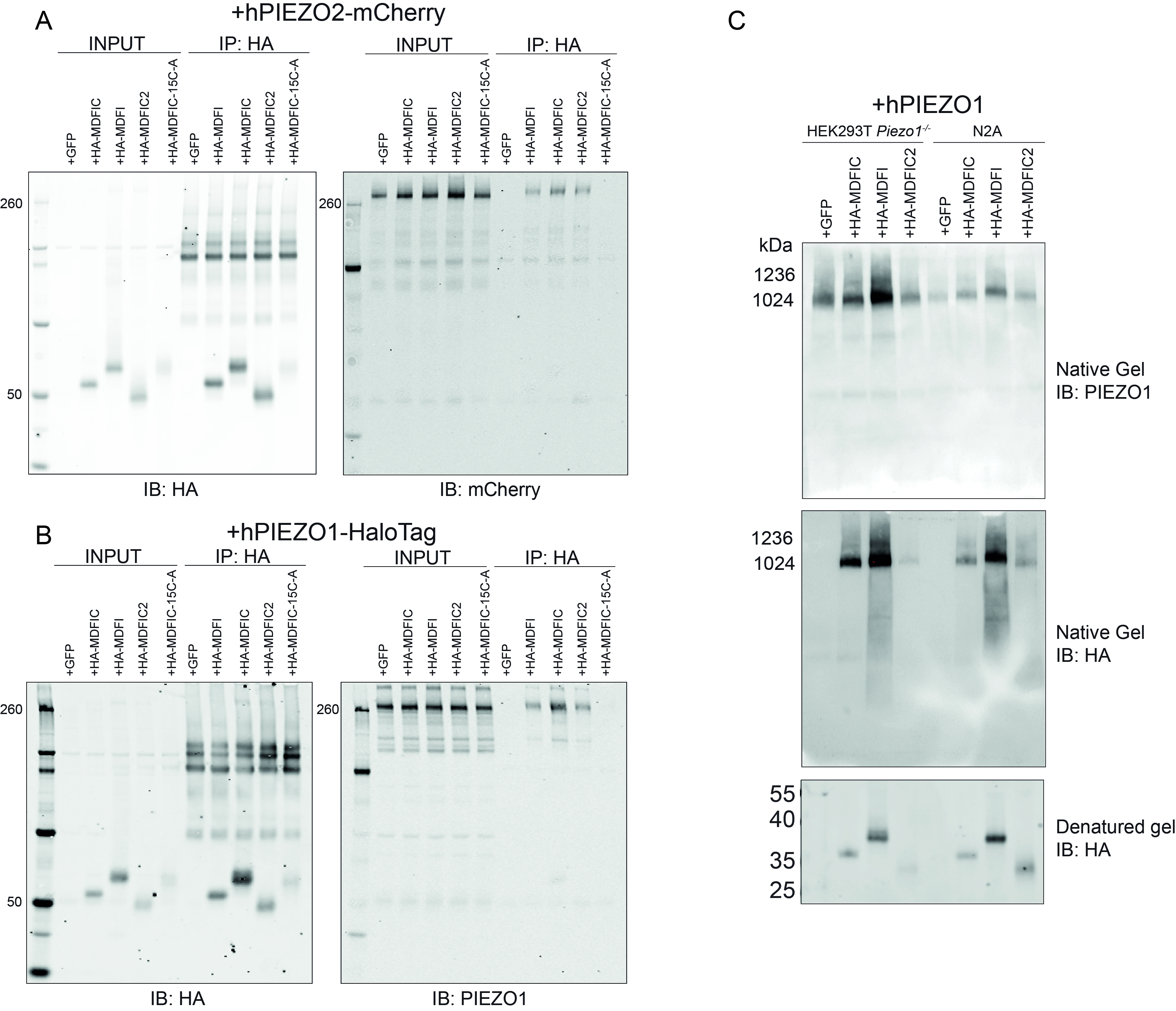

***Supplementary Figure 4. All MyoD-inhibitor family proteins bind PIEZO1 and PIEZO2.*** *All MyoD-inhibitor family proteins (MDFIC/MDFI/MDFIC2) could co-immunoprecipitate: (A) human PIEZO2-mCherry and (B) human PIEZO1-HaloTag. (C) Native gel of human PIEZO1 expressed with HA-MDFIC, HA-MDFI and HA-MDFIC2 with GFP as a vector only control in HEK293T cells and Neuro2A cells. Regardless of cell type we observed HA signal in the hPIEZO1 complexes for HA-MDFIC, HA-MDFI and HA-MDFIC2 using native gels. Top shows an anti-PIEZO1 immunoblot (NOVUS NBP2-75617), middle shows an anti-HA immunoblot indicating the presence of MyoD-inhibitor family proteins in the complex at the same size as the PIEZO1 trimer and the lower panel shows an SDS-page denaturing gel confirming the expression of HA-MDFIC, HA-MDFI and HA-MDFIC2 at the correct molecular sizes when denatured.*

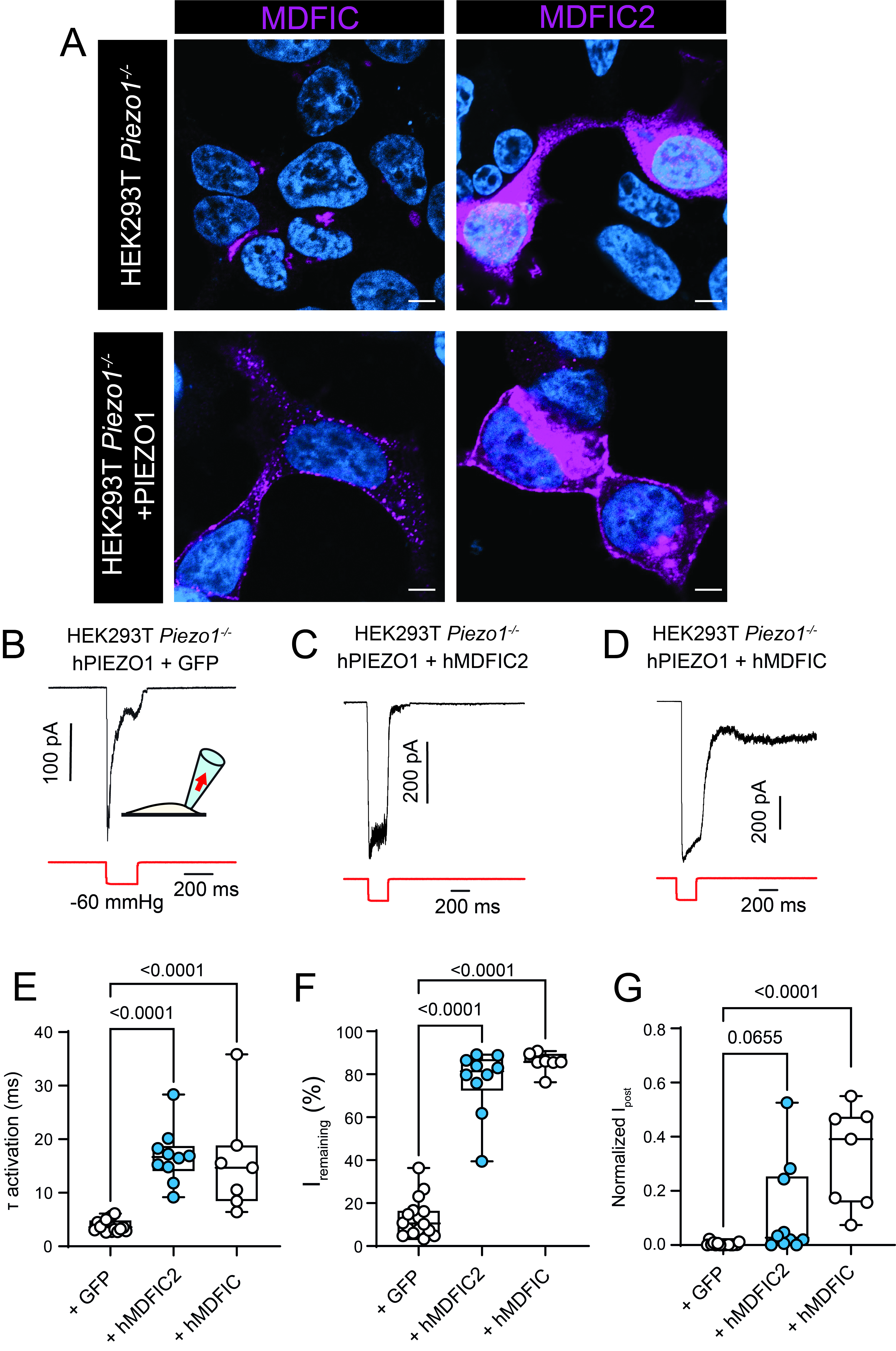

***Supplementary Figure 5. hMDFIC2 effects the biophysical properties of PIEZO1 in response to membrane stretch. (****A) Confocal images of HEK293T cells with and without PIEZO1 expressing HA-MDFIC or HA-MDFIC2 showing that on co-expression with human PIEZO1 hMDFIC2 is recruited to the membrane like hMDFIC. (B-D) Representative recordings in response to negative hydrostatic pressure (-60 mmHg) applied via a high-speed pressure clamp in the cell-attached configuration at a holding potential of -65 mV for PIEZO1 only, PIEZO1 plus hMDFIC2 and PIEZO1 plus hMDFIC for comparison. (E) The activation time constant of stretch-evoked currents is slowed by both hMDFIC and hMDFCI2 as measured by an exponential fit of current activation. (F) The current remaining (I_remaining_) at the end of the pressure pulse is increased by both hMDFIC2 and hMDFIC showing a profound reduction in inactivation. (G) The current still present 0.5 s after the end of the pressure pulse (I_post_) is affected only in the hMDFIC expressing cells not the hMDFIC2 expressing cells (Data is plotted as min to max box and whiskers plots; p-values shown determined using one way ANOVA with Tukey’s multiple comparison). (Scale bar – 5 μm)*

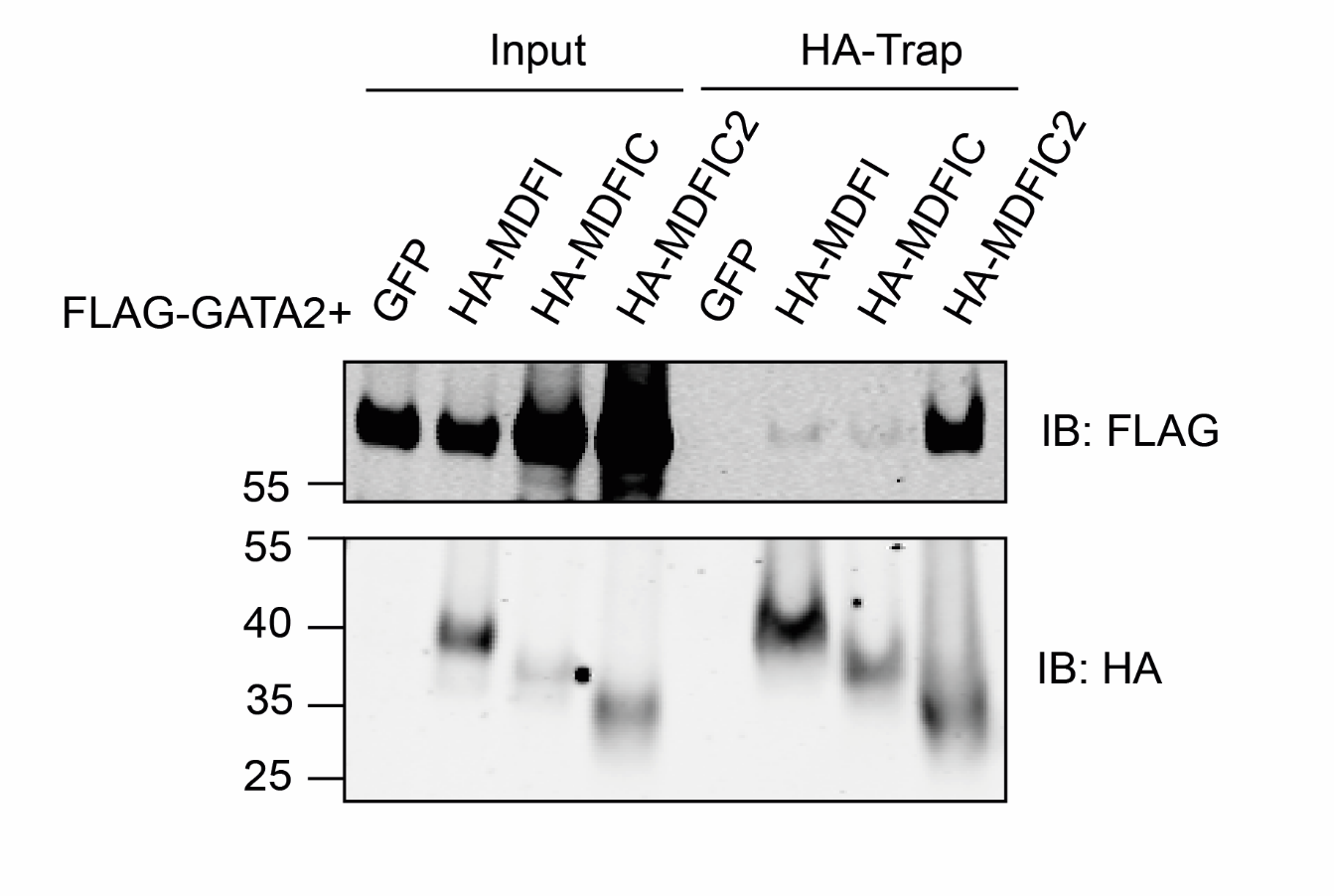

***Supplementary Figure 6. Co-immunoprecipitation shows all MyoD-inhibitor family proteins bind to GATA2.*** *Plasmids were transfected into Piezo1^-/-^ HEK293T cells and western blot shows on the left input fractions from cell lysate and on the right HA-tag immunoprecipitated fractions. The GATA2 was then probed with an anti-FLAG antibody and the MyoD-inhibitor family proteins* *were probed with an anti-HA antibody.*

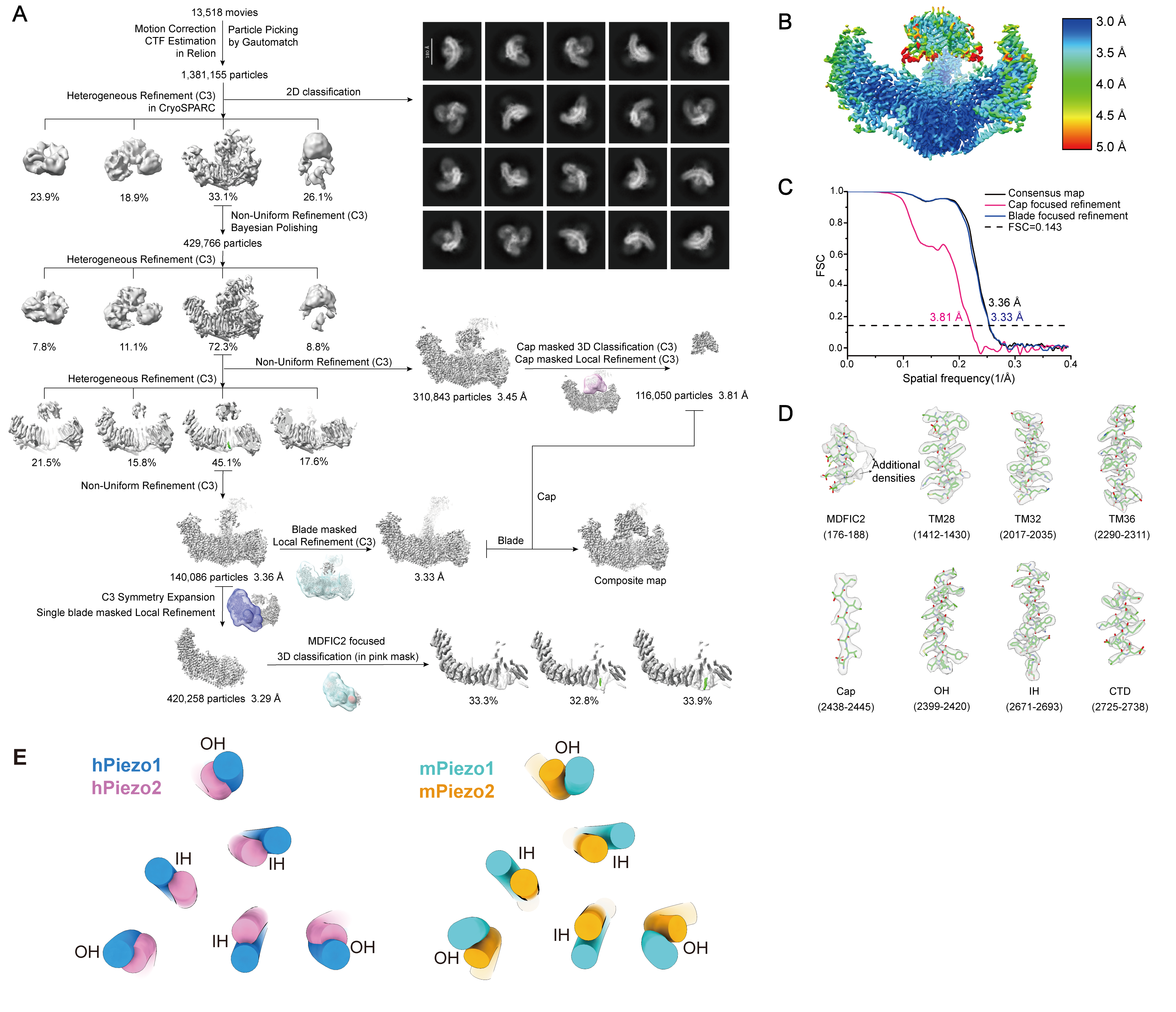

***Supplementary Figure 7. Cryo-EM data processing of the hPIEZO2-hMDFIC2 complex.*** *(A) Workflow of data processing from raw micrographs to high resolution structure determination. hMDFIC2 is colored green in the final round of heterogenous refinement and MDFIC2-masked 3D classification. (B) Cryo-EM map of the hPIEZO2-hMDFIC2 complex colored by local resolution. (C) Fourier shell correlation (FSC=0.143) curves calculated for the entire complex (black), the cap domain of hPIEZO2 (pink) and the body region (blue). (D) Representative cryo-EM densities of the hPIEZO2-hMDFIC2 complex. (E) Structural comparison of the pore module between PIEZO1 and PIEZO2 in human (left) and mouse (right). IH, inner helix; OH, outer helix. hPIEZO1, PDB:8ZU3; hPIEZO2, PDB:9VEE (this study); mPIEZO1, PDB:8IMZ; mPIEZO2, PDB:6KG7.*

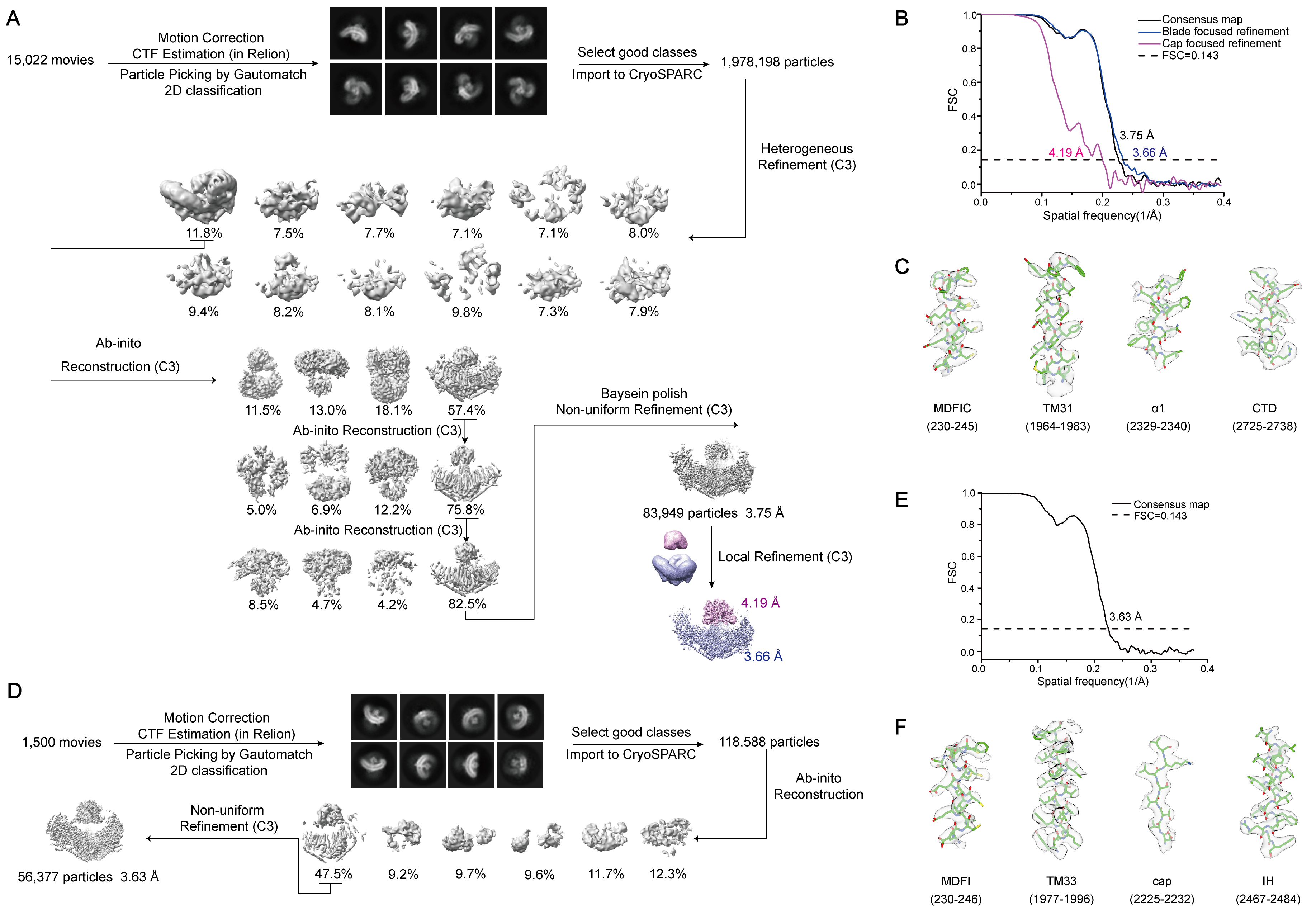

**Supplementary Figure 8. Cryo-EM data processing of the hPIEZO2-hMDFIC complex and mouse PIEZO1-mouse MDFI complex.** (A) Workflow of data processing from raw micrographs to high resolution structure determination of the hPIEZO2-hMDFIC complex. (B) FSC curves (FSC=0.143) calculated for the hPIEZO2-hMDFIC complex (black), the cap domain of hPIEZO2 (pink), and the body region (blue). (C) Representative cryo-EM densities of the hPIEZO2-hMDFIC complex. (D) Workflow of data processing from raw micrographs to high resolution structure determination of the mPIEZO1-mMDFI complex. (E) FSC curves (FSC=0.143) calculated for the mPIEZO1-mMDFI complex. (F) Representative cryo-EM densities of the mPIEZO1-mMDFI complex.

**Table S1. Cryo-EM Data collection, refinement and validation statistics**
This table provides comprehensive information on the Cryo-EM data collection, refinement, and validation statistics for the studied samples.

|  | hPIEZO2-hMDFIC2  composite map  (EMD-65005)  (PDB 9VEE) | hPIEZO2-hMDFIC2  consensus map  (EMD-64999)  (PDB 9VEE) | hPIEZO2-hMDFIC2  blade focused map  (EMD-65001)  (PDB 9VEE) | hPIEZO2-hMDFIC2  cap focused map  (EMD-65000)  (PDB 9VEE) | hPIEZO2-hMDFIC  composite map  (EMD-65006)  (PDB 9VEF) | hPIEZO2-hMDFIC  consensus map  (EMD-65002)  (PDB 9VEF) | hPIEZO2-hMDFIC  blade focused map  (EMD-65004)  (PDB 9VEF) | hPIEZO2-hMDFIC  cap focused map  (EMD-65003)  (PDB 9VEF) | mPIEZO1-mMDFI  (EMD-64998)  (PDB 9VED) |
| --- | --- | --- | --- | --- | --- | --- | --- | --- | --- |
| **Data collection and processing** |  |  |  |  |  |  |  |  |  |
| Magnification | 81,000 | 81,000 | 81,000 | 81,000 | 81,000 | 81,000 | 81,000 | 81,000 | 81,000 |
| Voltage (kV) | 300 | 300 | 300 | 300 | 300 | 300 | 300 | 300 | 300 |
| Electron exposure (e^-^/Å^2^) | 49.41 | 49.41 | 49.41 | 49.41 | 49.41 | 49.41 | 49.41 | 49.41 | 49.41 |
| Defocus range (μm) | -1.4 ~ -2.4 | -1.4 ~ -2.4 | -1.4 ~ -2.4 | -1.4 ~ -2.4 | -1.4 ~ -2.4 | -1.4 ~ -2.4 | -1.4 ~ -2.4 | -1.4 ~ -2.4 | -1.4 ~ -2.4 |
| Pixel size (Å) | 1.055 | 1.055 | 1.055 | 1.055 | 1.055 | 1.055 | 1.055 | 1.055 | 1.055 |
| Symmetry imposed | C3 | C3 | C3 | C3 | C3 | C3 | C3 | C3 | C3 |
| Initial particle images (no.) | 1,381,155 | 1,381,155 | 1,381,155 | 1,381,155 | 2,157,418 | 2,157,418 | 2,157,418 | 2,157,418 | 118,588 |
| Final particle images (no.) | 203,045 | 140,086 | 140,086 | 116,050 | 83,949 | 83,949 | 83,949 | 83,949 | 56,377 |
| Map resolution (Å) |  | 3.36 | 3.33 | 3.81 |  | 3.75 | 3.66 | 4.19 | 3.63 |
| FSC threshold |  | 0.143 | 0.143 | 0.143 |  | 0.143 | 0.143 | 0.143 | 0.143 |
| Map resolution range (Å) |  | 3.3-6.8 | 3.3-6.7 | 3.3-5.9 |  | 3.2-6.0 | 3.2-6.0 | 2.9-7.3 | 3.0-8.7 |
| **Refinement** |  |  |  |  |  |  |  |  |  |
| Initial model used (PDB code) | Predicted |  |  |  | Predicted |  |  |  | 8IMZ and predicted |
| Map sharpening *B* factor (Å^2^) |  | 111.1 | 105.2 | 142.3 |  | 123.2 | 105.9 | 189.3 | 116.1 |
| Model composition  Non-hydrogen atoms  Protein residues  Ligands | 34896  4251  0 |  |  |  | 34860  4254  0 |  |  |  | 29403  3732  0 |
| R.m.s. deviations  Bond lengths (Å)  Bond angles (°) | 0.009  0.891 |  |  |  | 0.010  0.912 |  |  |  | 0.012  0.998 |
| Validation  MolProbity score  Clash score  Poor rotamers (%) | 2.23  16.39  0.00 |  |  |  | 2.49  22.79  0.00 |  |  |  | 2.22  17.21  0.00 |
| Ramachandran plot  Outliers (%)  Allowed (%)  Favored (%) | 0.14  8.80  91.05 |  |  |  | 0.07  13.81  86.12 |  |  |  | 0.00  7.98  92.02 |

**Table S2. Antibodies used in this study.**

| **Designation** | **Source or reference** | **Catalogue number** | **Dilution** |
| --- | --- | --- | --- |
| anti-PIEZO1 | Novus Biological | NBP2-75617 | 1:1000 (WB)  0.6-1 µg (IP) |
| anti-MDFIC | Abcam | ab251808 | 1:200 (WB) |
| anti-HA | Cell Signaling | C29F4 | 1:1000 (WB)  66 ng (1 µl) (IP)  0.66 µg (10 µl) (IP-MS)  1:200 (IF) |
| anti-CD31 | BD Pharmigen | 550274 | 0.6-1 µg (EC isolation) |
| anti-Na/K-ATPase | DSHB | a6F | 1:1000 (WB) |
| anti-GAPDH | Cell Signaling | 2187 | 1:1000 (WB) |
| anti-mCherry (16D7) | Thermo Fisher Scientific | M11217 | 1:1000 (WB) |
| Anti-FLAG | Sigma | F3165 | 1:1000 (WB) |
| Goat anti-rabbit IgG conjugated to HRP | Abcam | ab6721 | 1:10000 (WB) |
| Goat anti-mouse IgG conjugated to IRDye 800CW | LICOR | 926-32210 | 1:10000 (WB) |
| Goat anti-rat IgG conjugated to HRP | Abcam | ab97057 | 1:10000 (WB) |
| Goat anti-rabbit IgG conjugated to IRDye 680RD | LICOR | 926-68071 | 1:10000 (WB) |
| Goat anti-rabbit conjugated to AF555 | Invitrogen | A21428 | 1:500 (IF) |

**Tables S3. Plasmid constructs used in this study.**

| **Recombinant DNA** | **Source** |
| --- | --- |
| Human PIEZO1-HaloTag in pIRES2 | (15) Addgene Plasmid #207834 |
| Human MDFIC pIRES2 | (Zhou et al., 2023) |
| Human HA-MDFIC pIRES2 | (Zhou et al., 2023) |
| Human HA-MDFIC2 pIRES2 | This study |
| Human HA-MDFIC2 Cys176/187Ala (2C-A) pIRES2 | This study |
| Human HA-MDFIC2 Δ20 (1-169) pIRES2 | This study |
| Human HA-MDFIC (15C-A) pIRES2 | (Zhou et al., 2023) |
| Human HA-MDFI in pIRES2 | (Zhou et al., 2023) |
| Mouse FLAG-MDFIC in pEG Bacmam | (Zhou et al., 2023) |
| Human FLAG-MDFIC pEG Bacmam | This study |
| Mouse FLAG-MDFI pEG Bacmam | This study |
| Human FLAG-MDFIC2 pEG Bacmam | This study |
| Human PIEZO2 pEG Bacmam | This study |
| Human PIEZO2-mCherry Codon optimized | This study fused to mCherry. Original clone from (16) |
| Human FLAG-GATA2 | Addgene Plasmid #1418 |
